# Model-free photon analysis of diffusion-based single-molecule FRET experiments

**DOI:** 10.1101/2024.10.31.621265

**Authors:** Ivan Terterov, Daniel Nettels, Tanya Lastiza-Male, Kim Bartels, Christian Loew, Rene Vancraenenbroeck, Itay Carmel, Gabriel Rosenblum, Hagen Hofmann

## Abstract

Photon-by-photon analysis tools for diffusion-based single-molecule Förster resonance energy transfer (smFRET) experiments often describe protein dynamics with Markov models. However, FRET efficiencies are only projections of the conformational space such that the measured dynamics can appear non-Markovian. Model-free methods to quantify FRET efficiency fluctuations would be desirable in this case. Here, we present such an approach. We determine FRET efficiency correlation functions free of artifacts from the finite length of photon trajectories or the diffusion of molecules through the confocal volume. We show that these functions capture the dynamics of proteins from micro-to milliseconds both in simulation and experiment, which provides a rigorous validation of current model-based analysis approaches.

---

Probing the dynamics of biomolecules has become a major task of single-molecule Förster resonance energy transfer (smFRET) experiments. Structures alone are insufficient to understand protein function. Instead, timescales and amplitudes of structural changes in enzymes^1-5^, transporters^6-8^, molecular machines^9,10^, and disordered proteins^11-14^ are required to understand their biological role. At the level of individual molecules, motions are stochastic and driven by thermal noise. Powerful tools to retrieve dynamics from stochastic trajectories are correlation functions. However, information in smFRET experiments, particularly with freely diffusing molecules, is scarce. Roughly 100 – 200 donor and acceptor photons are detected in a burst, i.e. during the short millisecond transit of a molecule through the confocal volume of a microscope. Although correlation functions of these short photon traces contain information on structural changes, it is obscured by the finite-length of the trajectory and by the diffusion process through the confocal volume itself. Therefore, innovative analysis tools to retrieve this information were developed with the goal to identify the number and type of structural states of a protein together with the timescales at which they are sampled. Established methods include dynamic PDA (photon distribution analysis)^15^, maximum likelihood (ML) methods^16-18^ and Hidden-Markov model fitting in the form of H^2^MM^1,19^ and mp-H^2^MM^20^. Despite differences in the details, these methods optimize the parameters of a model given the measured photon trajectory. Models are typically first-order chemical kinetic schemes of photon-emitting conformational states together with kinetic rate constants that describe switching between the states. To test the goodness of a fit, the photon traces are recolored using the model fit and then compared with the experimental data and/or the Viterbi algorithm is used to check whether the lifetimes of the states in the model are exponentially distributed^10^.

A drawback of these model-based approaches is that the model choice is not always obvious from the experimental FRET efficiency histograms. Several models need to be checked and a compromise between over-fitting and fit quality must be found. Often neglected, the FRET efficiency is a projection of the high-dimensional coordinate space of protein structures onto a single coordinate. Not all motions of a protein will necessarily cause a change in FRET efficiency, which can lead to apparent non-Markov behavior^21-23^ that might be missed by imposing Markov-models in the first place. An example is the enzyme QSOX (quiescin sulfhydryl oxidase) that samples two macroscopic structural states with power-law kinetics^2^. Model-free methods to extract dynamic information, e.g., in form of correlation functions, are therefore desirable. These functions provide the timescales of motions without necessarily imposing a model. In addition, they ease model identification for model-based analysis approaches and provide an additional test of the adequacy of a model. Current model-free methods to probe conformational dynamics in single-molecule bursts include lifetime-filtered FCS (fFCS)^24,25^, two-dimensional fluorescence lifetime correlation spectroscopy (2D-FLCS)^26-28^, time-resolved burst variance analysis (trBVA)^29^, and recurrence analysis of single particles (RASP)^13,30^. In these methods, the experimental photon traces are pre-processed to extract timescales of FRET efficiency fluctuations. Unfortunately, most of them require very long measurements to reach sufficient signal-to-noise (fFCS, 2D-FLCS, RASP) or they provide dynamic information in a rather indirect manner (trBVA).

Here, we present a simple but effective alternative. We show how to compute the autocorrelation function of FRET efficiencies^31^ free of artifacts due to the finite-length of the photon trace and the diffusion through the confocal volume. Using realistic simulations of smFRET experiments of diffusing molecules, we demonstrate that the timescale of FRET efficiency fluctuations can be correctly identified from microseconds up to milliseconds. Using experiments on a membrane protein and double-stranded DNA, we obtain dynamic information from as little as a few thousand molecules. We also show how this tool can be extended to probe the sub-microsecond dynamics of IDPs in nsFCS experiments^32-35^.

## Results

### Calculating FRET correlation functions

In smFRET experiments of freely diffusing molecules, the transit of a molecule through the confocal volume results in a burst of donor and acceptor photons (Fig. 1a). Unfortunately, a burst is only a few milliseconds long at best. If the burst duration ***T*** is close to the timescale of conformational dynamics (***τ***_***D***_), problems arise when attempting to extract ***τ***_***D***_ from correlation functions. The problem of computing correlation functions from continuous finite time series has been analyzed by Zwanzig^36^, who showed that the relative statistical error scales with 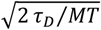 where ***M*** is the number of trajectories. An accuracy of 10% in the correlation function of a single molecule would require a burst that is 200 times longer than ***τ***_***D***_. With 200 trajectories on the other hand, even slow dynamics in the order of the burst duration (***τ***_***D***_**= *T***) might be accessible. Diffusion-based smFRET experiments with thousands of molecules should therefore be sufficient to determine correlation functions even for dynamics comparable to the burst duration. Yet, bursts are not continuous signals but rather streams of photons and their arrival times. We therefore distinguish three FRET efficiencies: the apparent FRET efficiency ***E***, defined by the raw photon counts of donor and acceptor, the FRET efficiency ***ϵ***, computed from the photon counts corrected for background, relative dye brightness and instrumental imperfections, and the continuous true FRET efficiency ***ε***, which is given by the time-dependent donor-acceptor distance ***r***(***t***) and the dye-specific Förster-distance ***R***_**0**_ (Supplementary Information eq. S1). To define experimental correlation functions, we assign an apparent FRET efficiency ***E* = 1** to each acceptor photon and ***E* = 0** to the donor photons. With ***τ*** being the time between two arbitrary photons in a burst, the FRET efficiency autocorrelation function (hereafter referred to as FRET correlation function) is formally given by

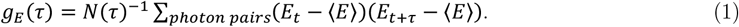

Here, ***N***(***τ***) is the number of all photon pairs separated by the lag time ***τ*** in all bursts and (***E***_***t***_, ***E***_***t*+*τ***_) indicates a specific photon pair separated by ***τ***. The sum is taken over all detected photon pairs separated by the lag time ***τ***. Importantly, **⟨*E*⟩** is the average FRET efficiency computed from the photons of all bursts. A proof that ***g***_***E***_(***τ***) **= *g***_***ε***_ (***τ***) at ideal instrumental conditions is provided in the Supplementary Information. Equation 1 can be reformulated (Supplementary Information). If ***N***_***XY***_(***τ***) is the number of photon pairs of type ***X*** and ***Y*** (***A*** for acceptor and ***D***for donor) separated by the time ***τ*** (Fig. 1b), we can write eq. 1 as

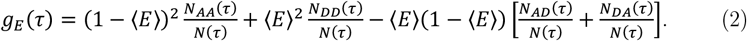

The fractions ***N***_***XY***_(***τ***)**⁄*N***(***τ***) are ratios of the conventional intensity correlation functions (Methods)

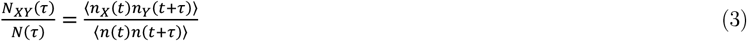

where ***n***_***X***_, ***n***_***Y***_ and ***n*** are the raw emission rates for photons of type ***X, Y***, and all photons, respectively. While ***N***_***XY***_(***τ***) and ***N***(***τ***) include intensity fluctuations due to the path of the molecule through the in-homogeneously illuminated confocal volume (Fig. 1c,d), these fluctuations cancel to large extends in eq. 3 (Supplementary Information). Similarly, bias due to the finite burst length^37^ is marginal (Supplementary Information). Hence, eq. 2 provides a good estimate of the FRET correlation function (Fig. 1d). Notably, a residual impact of diffusion on ***g***_***E***_(***τ***) will remain due to dye saturation and different detection volumes for donor and acceptor photons (Supplementary Fig. 1). However, while these effects indeed impact the ***N***_***XY***_(***τ***)**⁄*N***(***τ***) ratios, we show that they are negligible in ***g***_***E***_(***τ***) (see “Beyond the Poisson limit” and Supplementary Information).

**Fig. 1.**
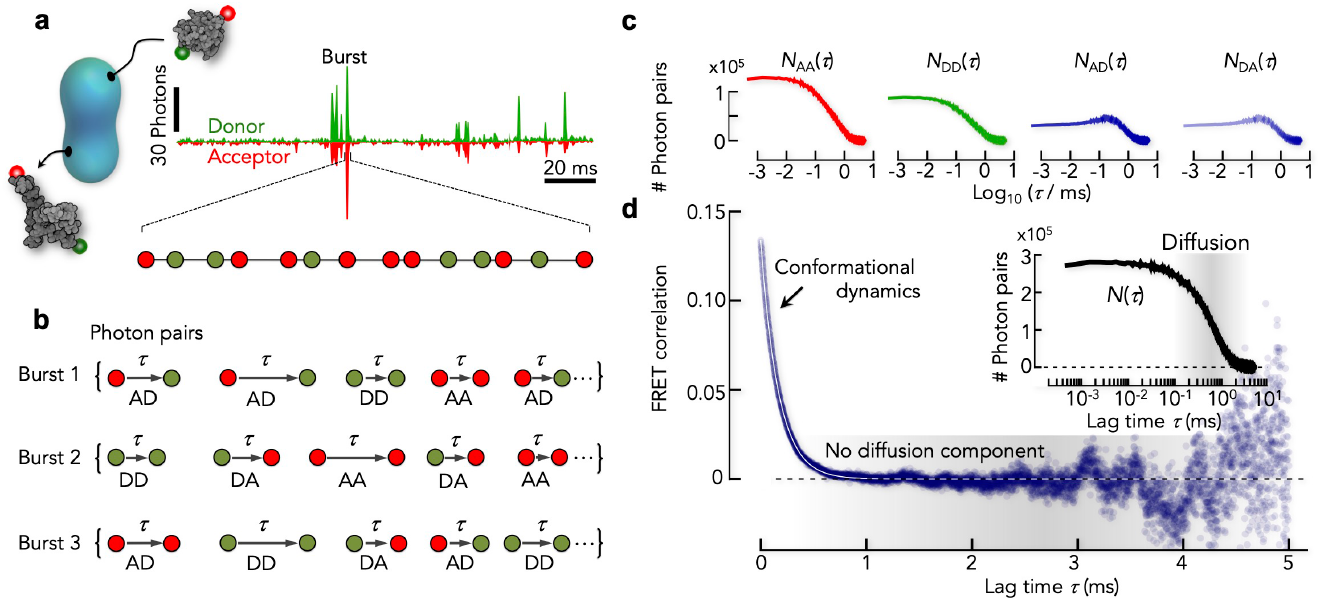
Calculating FRET correlation functions from photon trajectories. **(a)** The diffusion of donor-(D, green) and acceptor-(A, red) labeled proteins (gray) through the confocal volume (cyan, schematic) causes photon emission bursts of limited duration (*T*). **(b)** All possible pairs of photons together with their lag time (***τ***) are generated from the photon lists of all bursts. **(c)** Histograms of the lag times of each of the four types of photon pairs (***N***_***AA***_(***τ***), ***N***_***DD***_(***τ***), ***N***_***AD***_(***τ***), ***N***_***DA***_(***τ***)) are computed. **(d)** Dividing the photon pair histograms by the histogram of all photon pairs ***N***(***τ***) (inset), provides the correlation ratio ***N***_***XY***_(***τ***)**⁄*N***(***τ***). Using eq. 2, the FRET correlation function is computed (blue circles). The timescale of diffusion is indicated as gray shaded area.

### FRET correlation functions in the Poisson limit

To demonstrate the idea of our approach, we performed Brownian dynamics simulations of proteins at constant concentration that diffuse through a confocal volume and switch between two conformational states with the corrected FRET efficiencies ***ϵ***_**1**_ **= 0.1** and ***ϵ***_**2**_ **= 0.9** and the rate constants ***k* = *k***_**12**_ **= *k***_**21**_ (hereafter referred to as rates) (Fig. 2a). We added background photons, differences in the brightness of donor and acceptor, and the possibility to directly excite the acceptor by the donor excitation laser (Methods). The simulations assumed Poisson photon emission statistics that is correct at sufficiently low excitation rates and at timescales slower than the fluorescence lifetimes of the dyes. At low switching rates, the FRET histograms show two defined peaks (Fig. 2b). With increasing rates, intermediate FRET values become prominent because more molecules fchange their conformation during the transit through the confocal volume. At very fast exchange, the two peaks finally merge into a single peak at intermediate FRET efficiency, thus giving the impression of a single conformational state^38^. We then computed FRET correlation functions using eq. 2. These functions exhibit monotonic decays (Fig. 2c). The scatter in these decays increases with increasing lag-time due to the lower number of photon pairs that contribute to the correlation function at long times (Fig. 1c,d). Exponential fits describe the decays well (Fig. 2c). Given the 2-state model, the apparent relaxation rate (***λ***) is expected to be ***λ* = 2*k***. A comparison of the true relaxation rates with those determined from fits of the FRET correlation function shows excellent agreement (Fig. 2d). Notably, the correlation functions are exact, i.e., the value near zero lag time quantifies the variance of the FRET fluctuations. In our 2-state system, an approximate expression for the amplitude is ***g***_***E***_(**0**) **= *p***_**1**_***p***_**2**_(***E***_**2**_ **− *E***_**1**_)^**2**^ where ***p***_**1**_ **= *p***_**2**_ **= 1⁄2** are the relative populations of the two states and ***E***_**1**_, ***E***_**2**_ are the apparent FRET efficiencies of the two states (Supplementary Information). All functions decay to zero at long lag-times. However, static heterogeneity, e.g., due to a mixture of states that do not interconvert or that interconvert at timescales much slower than milliseconds, would manifest as an offset in the correlation functions. This is a particularly advantageous property. In fact, static heterogeneity is difficult to spot otherwise and is rarely included in model-based analysis approaches.

**Fig. 2.**
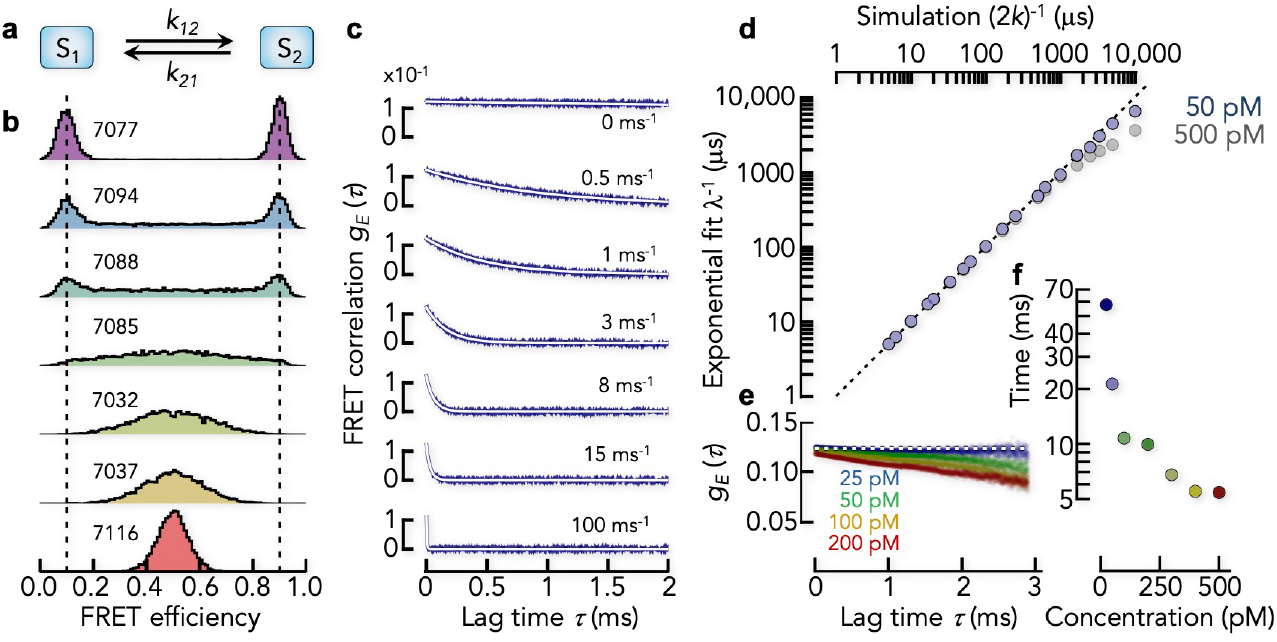
Determining dynamics from FRET correlation functions. **(a)** Kinetic scheme of the 2-state model. **(b)** FRET efficiency histograms (corrected) from Brownian dynamics simulations including the photon emission process of donor and acceptor. The FRET efficiencies of the two states in (a) are indicated by dashed lines. The total number of bursts is indicated for each histogram. The kinetic rates ***k* = *k***_**12**_ **= *k***_**21**_ (indicated in c) increase from top to bottom. **(c)** FRET correlation functions computed from the data in (b). The white line is a fit with an exponential decay and the apparent rate ***λ***. The kinetic rate used in the simulation is indicted. **(d)** Comparison between simulation and the analysis using FRET correlation functions. The total relaxation time from the exponential fits of the FRET correlation functions (***τ***_***app***_ **= 1⁄*λ***) is compared with the expected value ***τ***_***sim***_ **= 1⁄2*k***. The results of simulations with two protein concentrations (indicated) are shown. **(e)** FRET correlation functions for a static 2-state model (***k* = 0**) at different protein concentrations. The theoretical value of the FRET correlation function at infinite dilution is indicated by the dashed line. **(f)** Decay times of the FRET correlation functions in (e) after fits with an exponential decay.

### The effect of protein concentration on the correlation functions

Like in actual smFRET experiments, we simulated the data in Fig. 2a at the low concentration of 50 pM to ensure that the chance of simultaneously observing two or more molecules in the confocal volume is negligible. However, in some cases it is advantageous to perform smFRET experiments at higher concentrations, e.g., in inter-molecular FRET experiments to reach the affinity of the binding partners or if many photons are required such as in sub-population resolved nsFCS experiments^32,39,40^. Assuming a confocal volume of 1fl, Poisson statistics shows that 1.5% of the bursts include more than one molecule at a concentration of 50 pM. This fraction increases to 6% at 200 pM. These ‘mixed’ bursts unavoidably affect FRET correlation functions because two or more molecules in the confocal volume will average the FRET fluctuations^41^. We tested this effect by performing simulations at different protein concentrations. To exclusively study the concentration effect, we used a system in which two states with equal populations do not interconvert (static heterogeneity). At infinite dilution, the correlation function is expected to be a flat line at ***g***_***E***_(***τ***) **=** (***E***_**2**_ **− *E***_**1**_)^**2**^**⁄4** that quantifies the static heterogeneity of the mixture. Yet, with increasing protein concentration, ***g***_***E***_(***τ***) decays due to the de-correlation caused by multiple protein molecules in the confocal volume (Fig. 2e). The timescale of this decay is determined by the diffusion of the molecules and is rather long (>10 ms for concentrations <100 pM) (Fig. 2d,f). To avoid misinterpretations, we therefore suggest experiments at the lowest possible protein concentration for quantifying static heterogeneity or very slow dynamics. Notably, this concentration effect is inherent to the experiment and will affect every analysis method.

### Non-Markov processes and model validation

FRET efficiency fluctuations are 1D-projections of motions in a high-dimensional space spanned by the atomic coordinates of the protein. It can never be excluded that proteins explore states that are indistinguishable on the FRET efficiency coordinate. A recent example is Hsp90^42^. The dynamics of FRET efficiency fluctuations in this case will be non-Markovian with non-exponential dwell time distributions of the distinguishable states. Yet, analyzing dwell times first requires the assignment of states in the trajectory. While this is straightforward when photon fluxes are high, it is a daunting task in smFRET experiments of diffusing molecules. A few hundred photons per millisecond are insufficient to identify states without a model, which is the very idea behind model-based analysis tools. Yet, imposing Markov models on experimental data with potential non-Markov dynamics could be problematic. We demonstrate this aspect in a simulation. We simulated a 2-state model with FRET efficiencies ***ϵ***_**1**_ **= 0.2** and ***ϵ***_**2**_ **= 0.8** and the rates ***k* = 5 *ms***^**−1**^, which results in a broad FRET efficiency distribution centered between both states (Fig. 3a). A model-based analysis (see Methods) using the correct 2-state model provides an excellent fit as judged by a recoloring with the 2-state model and it reliably retrieves the simulated rates together with the FRET efficiencies of states 1 and 2. When using the recolored photon traces to compute the FRET correlation function, the simulated and fitted functions are in excellent agreement, as expected (Fig. 3b). Now, we render the system non-Markov by letting each state convert to a ‘mirror-image’ state (1’ and 2’) with the same FRET efficiency as the original state, leading to a 4-state model that appears 2-state on the FRET efficiency coordinate (Fig. 3c). For simplicity, we assume all rates to be identical (***k* = 5 *ms***^**−1**^). The simulation shows a broad distribution of FRET efficiencies with peaks at ***ϵ***_**1**_ **= 0.2** and ***ϵ***_**2**_ **= 0.8** and a distribution of bursts in between (Fig. 3c). Despite the increased complexity, a model-based analysis with a 2-state Markov-model provides an excellent fit of the data and there is no indication that the model might be inappropriate based on recoloring (Fig. 3c). The BIC (Bayes Information Criterion)^43^ is often used to determine the quality of a fit in Hidden-Markov modeling. It is given by ***BIC* = *R* ln *T*− 2ℒ**(***m***), where ***R*** is the number of free parameters in the fit, ***T***is the length of the trajectory, and **ℒ** is the maximized value of the log-likelihood function of the model ***m*** (see Methods). In our case (Markov vs. non-Markov), the ***BIC*** of fits with the 2-state-model to both data sets can only differ by **ℒ**. We found **ℒ = 16.8** for the fits of both data sets, thus providing no chance to distinguish the fit quality. Falsifying the model choice either requires a dwell-time analysis or additional model-independent information. Indeed, when we compared the FRET correlation function of the data with that obtained from the 2-state Markov-model fit, we found a substantial discrepancy (Fig. 3d). The correlation function of the data is non-exponential whereas the two-state fit results in a single-exponential decay. This comparison clearly invalidates the simple 2-state model in the case of non-Markov effects, thus calling for a more complex model.

**Fig. 3.**
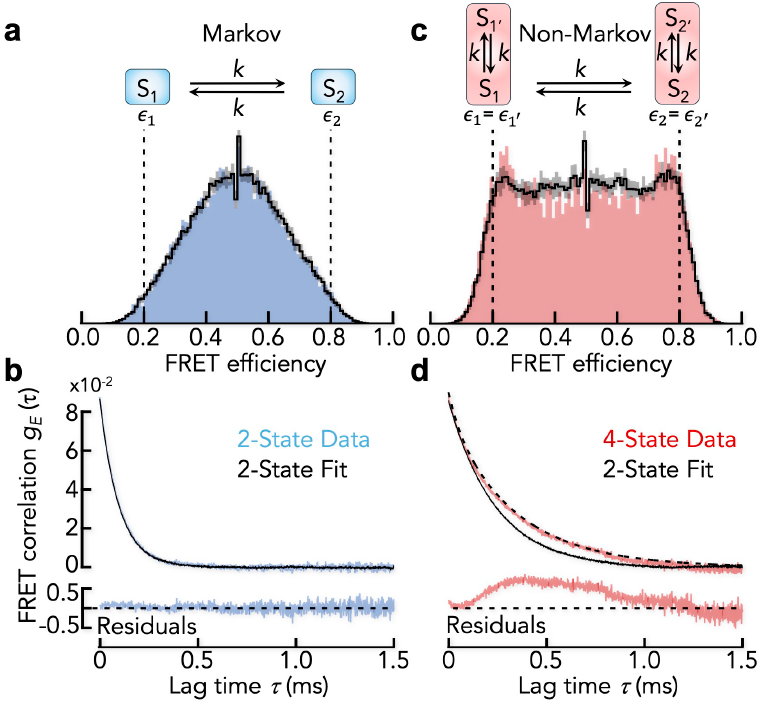
Model validation using FRET correlation functions. **(a)** FRET efficiency histogram (blue) of a simulation using a 2-state model (schematic) with interconversion rates k = 5 ms^-1^. Black line is the average of 10 recolored data sets based on a fit of the data with a 2-state Hidden-Markov model. The gray shaded area indicates 1 SD of the 10 realizations. The fitted rates are 5.1 ms^-1^ for the forward and backward reaction. **(b)** FRET correlation function (top) of the original data (blue) and the recolored data and the corresponding residuals of the fit (bottom). **(c)** FRET efficiency histogram (red) of a simulation using a 4-state model (schematic) with the rates k = 5 ms^-1^. Black line and gray shaded area as in (a). The fitted rates are 2.1 ms^-1^ for the forward and backward reaction. **(d)** FRET correlation function (top) of the original data (red) and the recolored data (black) and the corresponding residuals of the fit (bottom). The black dashed line is the analytically calculated FRET correlation function of the non-Markov model.

### Identifying photobleaching with correlation ratios^44^

The FRET correlation estimate can be decomposed into four correlation ratios (eq. 2) that fulfill the condition

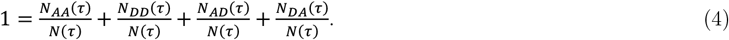

We define the normalized correlation ratios as

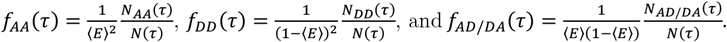

The correlation ratios contain similar information as ***g***_***E***_(***τ***) on the dynamics, but they are more prone to artifacts (see next section). Yet, they are helpful in identifying acceptor photobleaching due to chemical reactions of the dye with reactive oxygen species in the solution. To demonstrate this aspect, we performed simulations of a two-state system without and with acceptor photobleaching (Fig. 4a) and then computed the correlation ratios (Fig. 4b) and FRET correlation functions (Fig. 4c). In the absence of bleaching, the correlation ratios are symmetric with respect to a lag time of ***τ***= 0 ms (Fig. 4b left). Yet, when the acceptor of a molecule bleaches during the transit through the confocal volume, the density of donor photons will be higher towards the end of a burst than at its beginning. The cross-correlation ratios ***f***_***AD***_(***τ***) and ***f***_***DA***_(***τ***) will therefore not be identical (Fig. 4b middle). Acceptor-donor pairs (first acceptor then donor) are over-represented at long lag times whereas donor-acceptor pairs are underrepresented, which causes a pronounced asymmetry of the cross-correlation ratios. Similarly, donor photon pairs with long lag times are overrepresented, which causes a slow increase of ***f***_***DD***_(***τ***) with increasing ***τ*** paired with a decrease in ***f***_***AA***_(***τ***). When selecting only those bursts without acceptor bleaching using PIE (pulsed interleaved excitation)^45-47^ (Fig. 4a-b right), nearly symmetric cross-correlation ratios are retrieved. Hence, the cross-correlation ratios can be used to check whether bursts with photo-bleached acceptor dyes have been efficiently removed from the data. Notably, bleaching also affects the FRET correlation functions (Fig. 4c). Whereas appropriate filtering of only those bursts without bleached molecules retrieves the correct FRET correlation function (Fig. 4c right), the unfiltered data set causes an additional slow decay in the FRET correlation function (Fig. 4c middle), which is artificial.

**Fig. 4.**
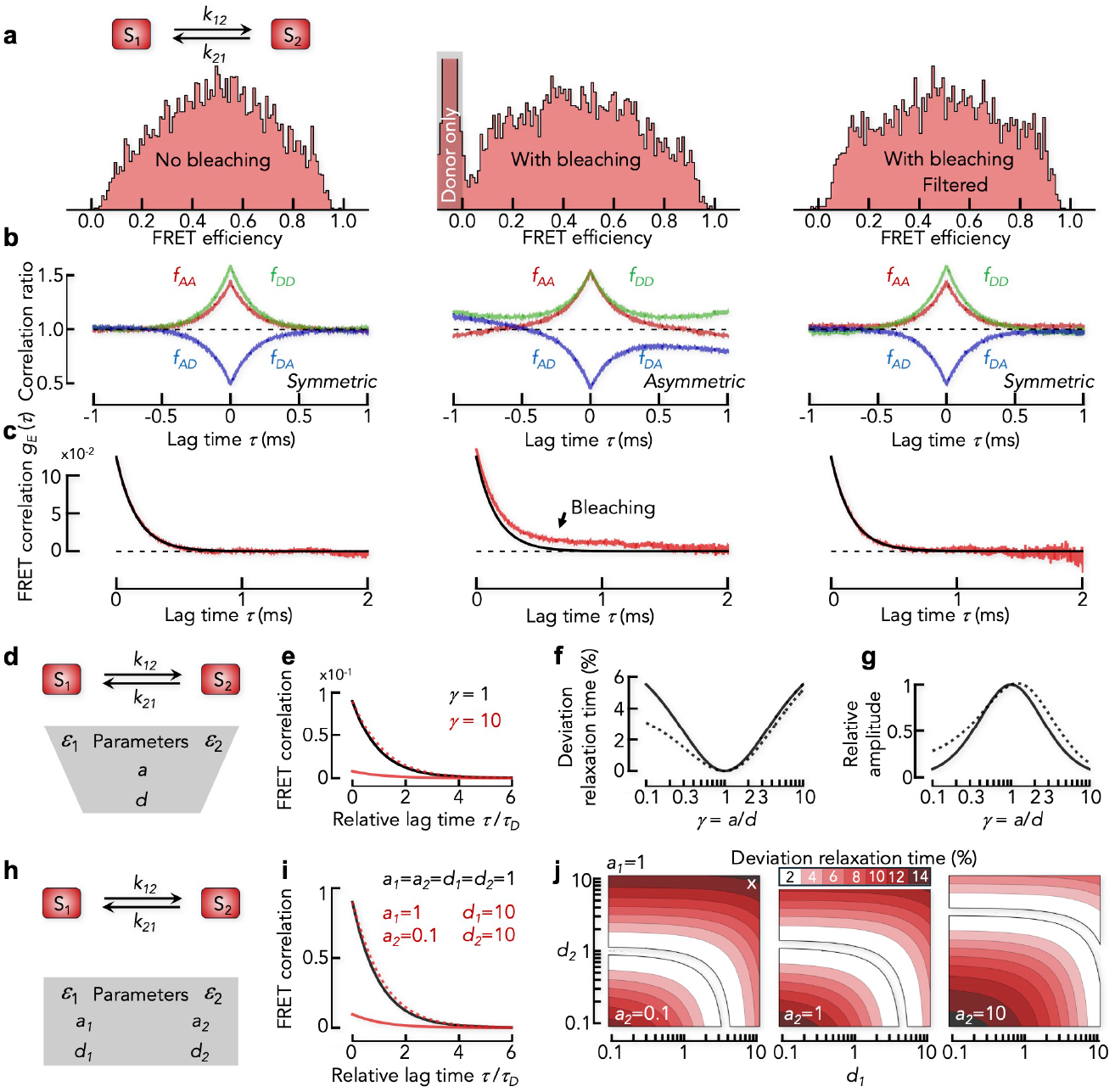
Impact of bleaching and dye brightness on FRET correlation functions. **(a)** Corrected FRET efficiency histograms for a two-state model (***k* = 3*ms***^**−1**^, ***ϵ***_**1**_ **= 0.1, *ϵ***_**2**_ **= 0.9**) without (left) and with (middle) bleaching. The gray shaded area indicates donor-only molecules that were excluded from the analysis (***ϵ* < 0** due to acceptor direct excitation). The FRET efficiency histogram with bleaching but filtered for unbleached bursts (right). **(b)** Normalized correlation ratios ***f***_***AA***_(***τ***) (red), ***f***_***DD***_(***τ***) (green), ***f***_***AD***_(***τ***) and ***f***_***DA***_(***τ***) (blue) for the data in (a). **(c)** FRET correlation for the data in (a). Black solid line is an exponential fit to the FRET correlation function in the absence of bleaching. **(d)** Kinetic scheme of the 2-state model with dye brightness independent of the conformational state. **(e)** Analytical solution of the FRET correlation function for the 2-state model for ***a* = *d*** (***γ* = 1**, black) and ***a* = 10*d*** (***γ* = 10**, red). The dashed line is the re-scaled correlation function shown for the case ***γ* = 10. (f)** Deviation of the relaxation rate from the true value without (solid black) and with background (dashed black). To highlight the difference, we used an unrealistic high background (10% of the acceptor signal). **(g)** Relative amplitude of the FRET correlation function. Solid and dashed lines are the same as in (f). **(h)** Kinetic scheme of the 2-state model with state-dependent dye brightness. **(i)** Analytical solutions of the FRET correlation function for identical brightness of all dyes and states (black line) and for a mixed case (red). The dashed red line is the re-scaled correlation function shown as solid red line. **(j)** Maps to indicate the relative deviation of the apparent relaxation rate (color scale) for the model in (h). Empty regions correspond to deviations <2%. A white cross indicates the case shown in (i).

### The impact of brightness differences on FRET correlation functions

The FRET correlation function ***g***_***E***_(***τ***) is computed from raw photon traces. Experimental imperfections such as differences in the quantum yields of the dyes (***Q***_***A***_, ***Q***_***D***_) or efficiencies of the detectors (**ζ**_***A***_, **ζ**_***D***_) render ***g***_***E***_(***τ***) different from the true FRET correlation function ***g***_***ϵ***_(***τ***). The procedures to account for these imperfections in the calculation of correct FRET efficiencies (***ϵ***) have been discussed extensively in the past^46,48,49^. Let ***n***_**0**_ be the photon emission rate of each dye at identical excitation rates in the absence of imperfections, then ***a* = ζ**_***A***_***Q***_***A***_***n***_**0**_ and ***d* = ζ**_***D***_***Q***_***D***_***n***_**0**_ are the measured photon rates (brightness) of acceptor and donor in a microscope. The correction factor ***γ* = *a*⁄*d*** is typically used to compute corrected FRET efficiencies. For a 2-state system with brightness differences between the dyes, the measured and true FRET correlation functions are related by (Methods)

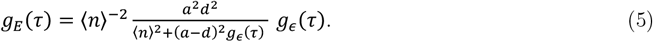

Here, **⟨*n*⟩ = *a*⟨*ϵ*⟩ + *d***(**1 − ⟨*ϵ*⟩**) is the total photon rate averaged over the conformational states. The factor in front of ***g***_***ϵ***_(***τ***) is time-dependent due to ***g***_***ϵ***_(***τ***) in the denominator, which alters the decay of ***g***_***E***_(***τ***) compared to ***g***_***ϵ***_(***τ***) (Fig. 4d,e). Yet, its impact is marginal because **⟨*n*⟩**^**2**^ dominates the denominator. Indeed, within the range **0.1 ≤ *γ* ≤ 10**, which by far exceeds correction factors in most smFRET experiments, the relaxation time of a 2-state system obtained from ***g***_***E***_(***τ***) does not deviate more than 6% from the true value (Fig. 4f). This deviation is further diminished in the presence of background photons. Contrary to the relaxation time, the amplitude of ***g***_***E***_(***τ***) is strongly affected by differences in the dye brightness (Fig. 4g). The timescales determined with FRET correlation functions on the other hand, are rather robust.

Yet, the dye brightness might be a fluctuating quantity, and two cases can be distinguished: the brightness is dependent (case 1) or independent (case 2) of the conformational states. In case 1, the result depends on the kinetic model (Fig. 4h). For a 2-state model, the FRET correlation function is given by

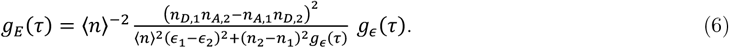

Here, the total photon rate of state ***i*** is ***n***_***i***_ **= *n***_***A***,***i***_**+*n***_***D***,***i***_ with ***n***_***A***,***i***_ **= *a***_***i***_***ϵ***_***i***_ and ***n***_***D***,***i***_ **= *d***_***i***_(**1 − *ϵ***_***i***_) and the average total photon rate is **⟨*n*⟩ = *p***_**1**_***n***_**1**_ **+ *p***_**2**_***n***_**2**_. Again, the leading term in the denominator (**∝ ⟨*n*⟩**^**2**^) is time-independent such that ***g***_***E***_(***τ***) decays like the true correlation function ***g***_***ϵ***_(***τ***) (Fig. 4i). Indeed, calculations of a 2-state model show that the relaxation time differs not more than 16% from the true value even if ***a***_***i***_ and ***d***_***i***_ vary by two orders of magnitude (Fig. 4j).

If the dye brightness fluctuates independently of the structural state of the system, ***g***_***E***_(***τ***) is a combination of the photophysical and structural dynamics whose behavior depends on the specific processes that lead to the brightness fluctuations. General conclusions are difficult to draw in this case unless the fluctuations are much faster than the conformational dynamics in which case eq. 5 will be obtained. In summary, brightness differences of the dyes impact FRET correlation functions. Yet, within experimental limits, the error in timescale is marginal unless the brightness fluctuates independently of conformational transitions. Like model-based analysis approaches, a correlation analysis quantifies fluctuations in raw FRET efficiencies, but it does not provide the means to ambiguously identify the source of these fluctuations.

### Beyond the Poisson limit

So far, we modeled the emission of photons as a Poisson process because the nanosecond lifetimes of photophysical singlet states of the dyes are much faster than the microsecond dynamics we are interested in. However, organic fluorophores are large conjugated *π*-systems that also populate triplet states with microsecond lifetimes. Probably best characterized is the dye pair AlexaFluor488 and AlexaFluor594 (Fig. 5a)^50^. Nettels *et al*. determined the photophysical model for this pair including the transition rates (Supplementary Table 1) and showed that the FRET efficiency depends weakly on the laser power^50^. Albeit this dependence is less relevant for FRET histograms due to singlet-singlet and singlet-triplet annihilation processes^50^, it might impact the dynamics extracted from diffusion-based smFRET experiments. When a molecule diffuses through the confocal volume, excitation intensity, photon emission rates, and ultimately also the measured FRET efficiency are functions of the position in the confocal volume. To estimate the magnitude of these non-idealities, we performed Brownian dynamics simulations of molecules with a donor-acceptor distance fixed to the value of the Förster distance (Fig. 5b). Electronic transitions take place at two timescales: nanoseconds for transitions out of singlet states and microseconds for transitions out of triplet states. We take advantage of this timescale separation and coarse-grained the photophysical scheme such that the dynamics at microsecond timescales is approximately preserved in the simulation (Fig. 5a, see Methods and Supplementary Information). An overlay of the simulated photon traces based on their mean photon arrival times indeed shows a mismatch of donor and acceptor photon rates at the center of the confocal volume where the excitation intensity is highest (Fig. 5c). This results in a time-dependent FRET efficiency due to the diffusion of molecules through the confocal volume (Fig. 5d). This combination of triplet dynamics and illumination-induced FRET fluctuations strongly affects intensity correlation functions^51^ and correlation ratios. We find that ***f***_***AA***_(***τ***) first decreases and then increases again (Fig. 5e and inset). The donor ratio ***f***_***DD***_(***τ***) shows the same decays albeit with reversed amplitude signs, i.e., ***f***_***DD***_(***τ***) first increases due to triplet dynamics and then slowly decreases due to FRET fluctuations (Fig. 5e and inset). The cross-correlation ratios (***f***_***AD***_(***τ***) and ***f***_***DA***_(***τ***)) are less affected, but they also show a fast and slow component due to triplet dynamics and diffusion, respectively. Notably, these components are significantly suppressed in the FRET correlation function (Fig. 5f). In fact, the slow component is virtually absent because the FRET-changes when passing the excitation volume are exceedingly small (Fig. 5d). However, the fast decay resulting from the lifetime of triplet states is observed at a timescale of 3 μs. Although the triplet amplitude is predicted to be small (Fig. 5f), care should be taken in model-based analysis approaches that do not explicitly include these dye dynamics. As a complete modeling of photophysical transitions is nearly impossible given the incomplete characterization of organic fluorophores used in smFRET experiments, we recommend an empirical approach to minimize the impact of triplet dynamics. For instance, triplet dynamics can be reduced at lower laser powers, choosing dyes with low triplet occupancy, or by using triplet-quenching additives.

**Fig. 5.**
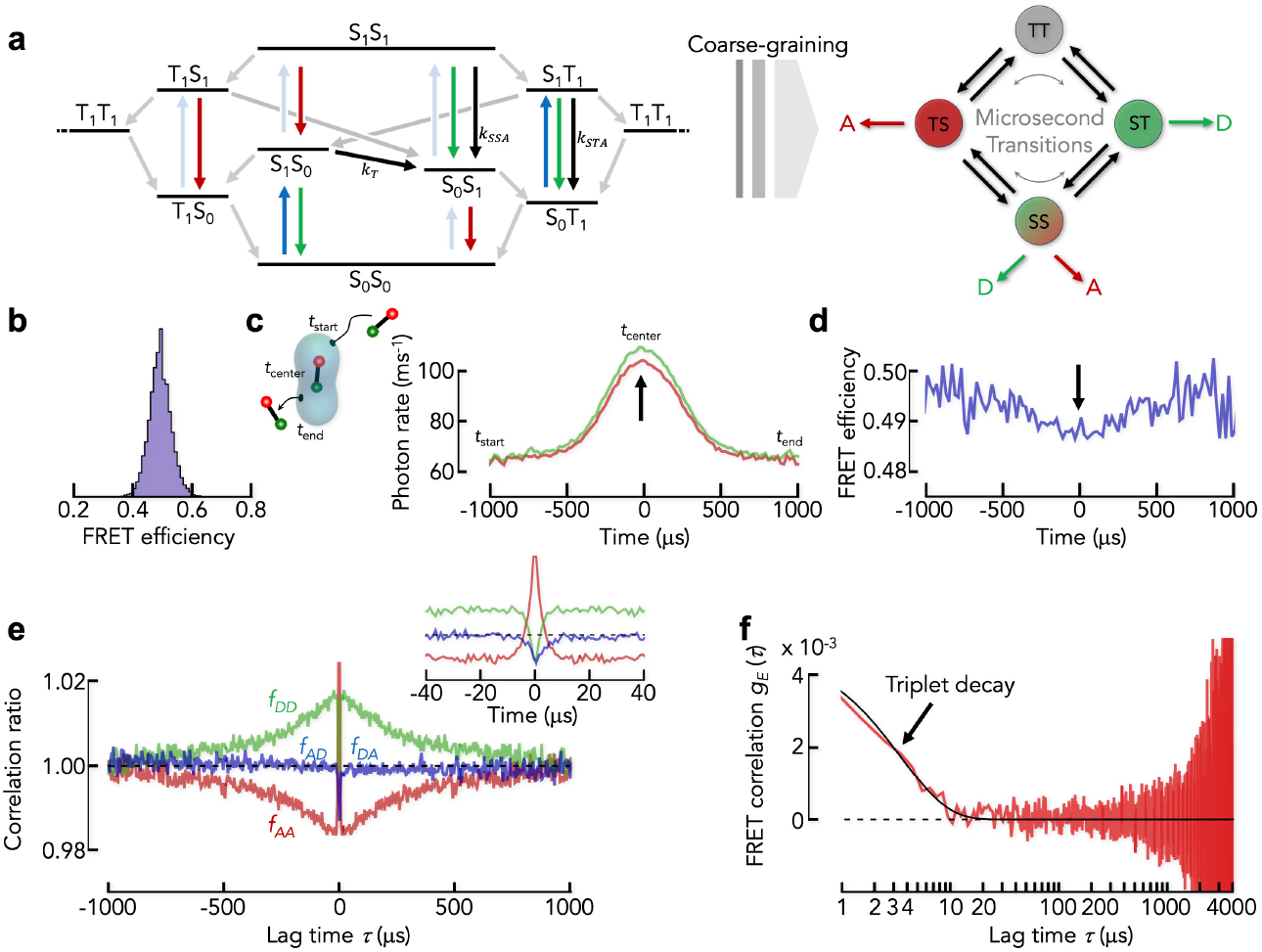
The impact of non-Poissonian photon emission. **(a)** Jablonski diagram of the dye pair AlexaFluor 488 and 594 based on Nettels *et al*.^50^ (left). Each state is denoted by the electronic states of donor (first symbol) and acceptor (second symbol). S and T stand for the singlet and triplet manifolds, respectively. Subscripts refer to electronical ground (0) and first excited (1) states. The first letter refers to the donor and the second letter indicates the state of the acceptor. Red and green arrows indicate transitions that lead to the emission of acceptor and donor photons, respectively. Dark blue arrows are excitation transitions of the donor and light blue arrows indicate the direct excitation of the acceptor at the wavelength of the donor. Black arrows indicate energy transfer processes where singlet-singlet annihilation and singlet-triplet annihilation are indicated by the rates ***k***_***SSA***_ and ***k***_***STA***_, respectively (Supplementary Table S1). The classical energy transfer rate is ***k***_***T***_. Gray arrows indicate singlet-triplet and triplet-singlet transitions. The coarse-grained (CG) model of the full photophysical scheme (right) has 4 states that interconvert at microsecond timescales. **(b)** FRET histogram of a particle with fixed donor-acceptor distance identical to the Förster distance simulated using the coarse-grained model in (a). **(c)** Average of all photon traces of the bursts in the simulation. The overlay was constructed by aligning the trajectories relative to the average arrival time of donor (green) and acceptor (red) photons. The mean arrival time was arbitrarily set to zero. Arrow indicates the mismatch between donor and acceptor signal. **(d)** FRET efficiency profile calculated from the data in (c). **(e)** Normalized correlation ratios for the data in b-d. Inset: Zoom of the normalized correlation ratios. **(f)** FRET correlation function for the data in b-d. Black line is a single-exponential fit. Arrow indicates the triplet-induced decay.

### Experimental applications

To demonstrate the application of FRET correlation functions, we investigated the dynamics of (i) a membrane protein proton dependent oligo-peptide transporter (POT), (ii) double-stranded DNA (dsDNA), and (iii) an intrinsically disordered protein (a modified DNA-binding domain of c-Myc).

The bacterial di- and tripeptide permease DtpA (*E. coli*) (Fig. 6a, inset) uses a proton gradient to transport di- and tri-peptides across the membrane^52^. The two helix bundles of DtpA have been proposed to open and close in an alternating fashion. Substrate and protons bind at the periplasmic side of the protein. A conformational change then closes the periplasmic side and opens the cytoplasmic side, thus releasing them. Using RASP^30^, we showed previously that the distance between the helix bundles on the cytoplasmic side fluctuate at a timescale of ∼1 ms when the protein is embedded in LMNG micelles^8^. To confirm these dynamics, we used the DtpA-variant W203C/Q487C with donor (AlexaFluor488) and acceptor (AlexaFluor594) being attached to the helix bundles at the cytoplasmic side (Fig. 6a, inset). The FRET histogram of the transporter shows a broad distribution that indicates a heterogeneous mixture of different conformational states (Fig. 6a). The correlation ratios resemble the pattern found in our simulations using the coarse-grained photophysical scheme (Fig. 5e), which demonstrates that photophysical non-idealities such as triplet dynamics and the dependence of FRET efficiency on the laser excitation are indeed found in experimental data (Fig. 6b). The acceptor autocorrelation ratio first decreases and then increases slightly whereas the donor autocorrelation ratio first increases and then decreases (Fig. 6b). The cross-correlation ratios increase monotonically and are symmetric for DA and AD photon pairs, suggesting that bursts with bleached acceptor had been successfully removed from the data. The FRET correlation function shows a complex decay with multiple components and an offset (Fig. 6c). In lack of a physical model for the dynamics of DtpA, we empirically fitted the data with a sum of up to five exponential decays and a manually determined offset to determine an empirical relaxation time (Fig. 6d). We recommend determining offsets manually based on the data points with the longest lag time to obtain an upper estimate of the static heterogeneity. An inspection of the residuals of the fits indicates that three exponential decays are sufficient to describe the data set. The relaxation times are ***τ***_1_ = 8 μs,***τ***_2_ = 117 μs, and ***τ***_3_ = 1.8 ms. The fastest timescale coincides with triplet dynamics (Fig. 6c, 5f). However, the amplitude of this fast decay is substantially higher than in our simulations (Fig. 5f), which either indicates that the photophysical model is incomplete or that protein motions mix with triplet dynamics. Importantly, the slower processes are within the range found with RASP. In fact, a weighted average of the slow relaxation times gives ∼1.2 ms, in good accord with the value found with RASP (∼1 ms)^8^. Also, the offset indicates the presence of motions slower than milliseconds as found with RASP^8^.

**Fig. 6.**
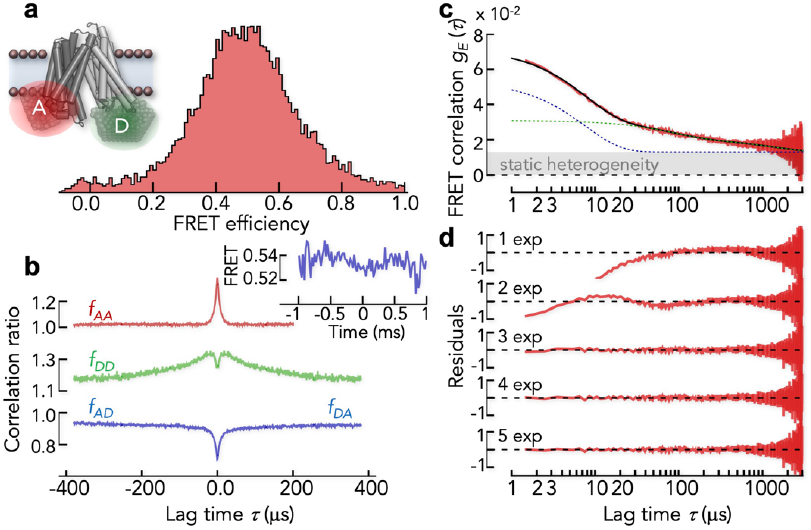
Dynamics of the membrane protein DtpA. **(a)** FRET efficiency histogram of DtpA in the detergent LMNG labeled at the cytoplasmic side of the two helix bundles (inset). **(b)** Normalized correlation ratios of DtpA and averaged FRET trajectory of all bursts (inset). **(c)** FRET correlation function of DtpA computed from the data in (a). Solid black line is a fit with a sum of three exponential decays. Dashed blue line is the decay of the fastest component and green dashed line is the sum of all other decay components. The gray shaded area indicates the static heterogeneity (offset). **(d)** Residuals of the fits with multiple exponential decays. The number of exponentials is indicated.

In a second example, we investigated the dynamics of dsDNA. Early FCS experiments identified complex structural dynamics at microsecond timescales^53^. We previously constructed a set of 12 dsDNAs labeled with donor (AlexaFluor 488) and acceptor (AlexaFluor 594) dyes^29,54^. The experiments were performed at neutral (pH 7) and acidic (pH 4) conditions since DNA is destabilized at low pH^55^, which alters the dynamics^29^. The FRET efficiency histograms of the 12 samples span the complete FRET range (Fig. 7a) and the correlation functions show a monotonic decay for all of them (Fig. 7b). Overall, we found higher amplitudes at pH 4, which reflects the increased flexibility of dsDNA at destabilizing acidic conditions^55^. Similarly to DtpA, we globally fitted the decays at each pH using a sum of exponentials with manually defined offsets to account for static heterogeneity. The change in the fit quality indicated that three exponentials were again sufficient to describe all correlation functions (Fig. 7c) with the fastest timescale (5 μs at pH 7, 10 μs at pH 4) being close to the triplet timescale. The intermediate timescale (***τ***_2_ = 72 μs at pH 7, ***τ***_2_ = 62 μs at pH 4) is in accord with the 50 μs found previously for the dynamics of dsDNA. Yet, the amplitudes of the slowest decay (305 μs at pH 7, 405 μs at pH 4) increase with increasing mean FRET values of the samples (Fig. 7b,d). This long timescale has previously been interpreted as a slow mode of DNA motion based on the rather indirect trBVA analysis^29^. Our analysis now provides additional diagnostic tools to falsify this conclusion. In particular, the incomplete removal of bursts with a bleached acceptor can cause such a slow decay in the FRET correlation function (Fig. 4c, middle). An asymmetry of the cross-correlation ratios ***f***_***AD***_(***τ***) and ***f***_***DA***_(***τ***) easily identifies such effects (Fig. 4b, middle). Indeed, we found that the cross-correlation ratios are symmetric for samples with low FRET efficiency, thus ruling out acceptor bleaching (Fig. 7e *left*). At high FRET efficiencies on the other hand, the acceptor resides more frequently in the excited state, thus increasing the chance for photo-oxidation. We find that the cross-correlation ratios are highly asymmetric for these samples (Fig. 7e *right*), suggesting that acceptor bleaching is indeed the main contribution to the slow decay in the FRET correlation functions. Albeit fast (< 10 μs) and very slow (> 100 μs) structural dynamics are hard to exclude with absolute certainty, our FRET correlation analysis confirms the previously found timescale of 50 μs.

**Fig. 7.**
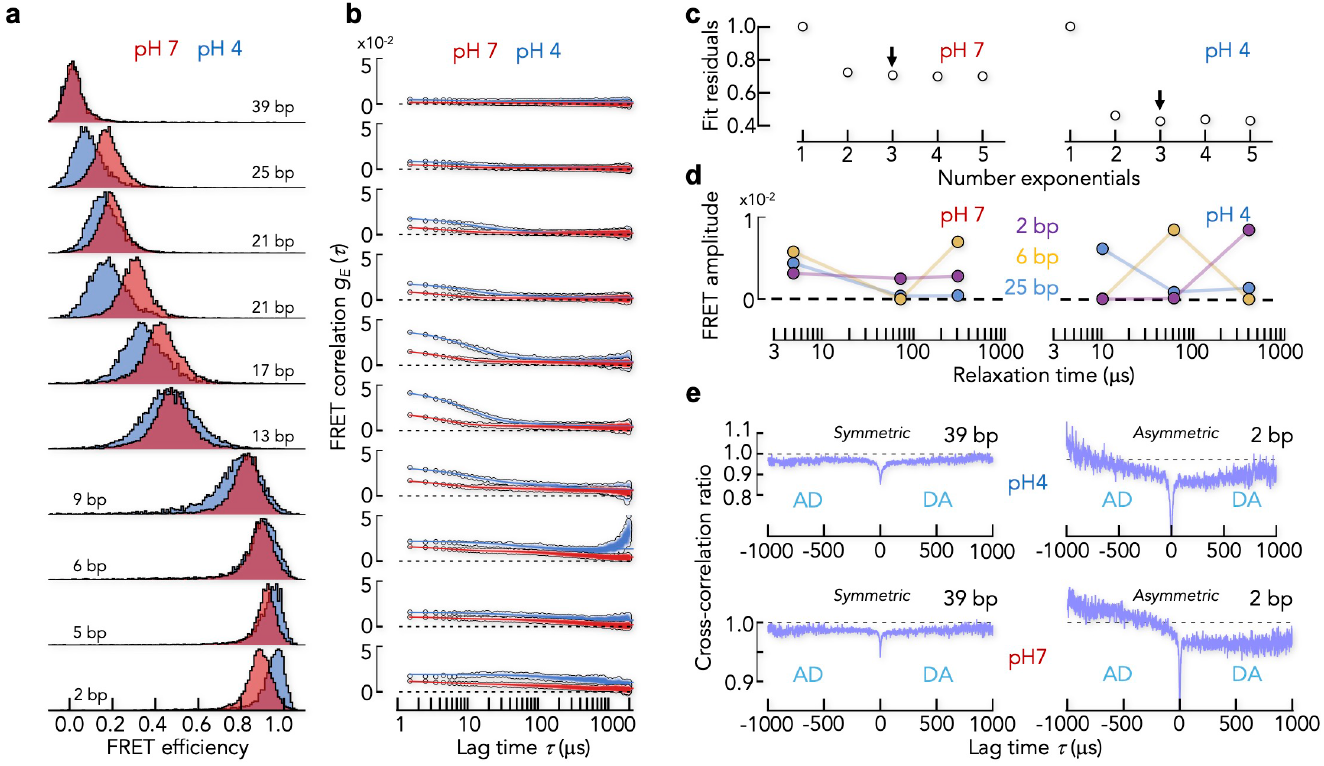
Dynamics of dsDNA probed with FRET correlation functions. **(a)** FRET histograms of dsDNA samples with decreasing sequence separation (indicated) at pH7 (red) and pH4 (blue). **(b)** FRET correlation functions for the data in (a). Solid lines are global fits with a sum of three exponential decays and an offset. **(c)** Summed fit residuals for fits with different numbers of exponential decays for the data and fits at pH7 (left) and pH4 (right). **(d)** Amplitudes of the three decays plotted as function of the determined relaxation times for three sequence separations for pH7 (left) and pH4 (right). **(e)** Normalized cross-correlation ratios at pH4 (top) and pH7 (bottom) for high sequence separation, i.e., low FRET (left), and low sequence separation, i.e., high FRET (right).

Finally, we demonstrate that FRET correlation functions are not restricted to microsecond timescales. We investigated the IDP ΔMyc, which is a modified version of the DNA-binding domain of the transcription factor c-Myc. In this modification, all hydrophobic residues were replaced by serine and glycine residues^56^, which ensures that this sequence is largely unstructured. We recently studied the reconfiguration dynamics of ΔMyc^35^ and found fast sub-microsecond timescales as expected for an IDP^11,32,33,39,57-59^. We therefore do not expect dynamics at timescales of microseconds. We performed sub-population specific nsFCS experiments^39^ with ΔMyc at a denaturant concentration of 2 M GdmCl to exclude any residual transient structure formation. We collected > 800,000 bursts to also probe the nanosecond dynamics with sufficient photon pair statistics (Fig. 8a, inset). The classical intensity correlation functions show five decay components: Anti-bunching, conformational dynamics, triplet dynamics, an unknown component, and the decay due to the diffusion of molecules through the confocal volume (Fig. 8a). Conformational dynamics due to chain reconfigurations differ from other decays by a rise in the cross-correlation functions. Compared to the complexity of the intensity correlation functions, the FRET correlation function is simple (Fig. 8b). Only antibunching, reconfiguration and triplet dynamics are left with significant amplitude whereas the dominant diffusion component is efficiently suppressed. The reconfiguration timescales are 38 ns (intensity correlation) and 43 ns (FRET correlation), i.e., very similar. The slower timescale of triplet dynamics agrees well with our simulations of the photophysical scheme both in terms of timescale (3 μs) and amplitude (0.006) (see Fig. 5f for comparison). The un-assigned decay at 27 μs that is prominent in the intensity correlation functions, is also substantially suppressed in the FRET correlation function, indicating that it does not result from changes in FRET efficiency. Finally, due to the lack of dynamics at timescales >30 μs, the data provide a chance to check the quality of diffusion suppression in FRET correlations. A zoom at microsecond timescales shows residual fluctuations with an amplitude of 0.002 (3% of the total amplitude). This value can be considered as a lower limit, i.e., the amplitude of decays in FRET correlation functions should be > 0.002 to be considered significant.

**Fig. 8.**
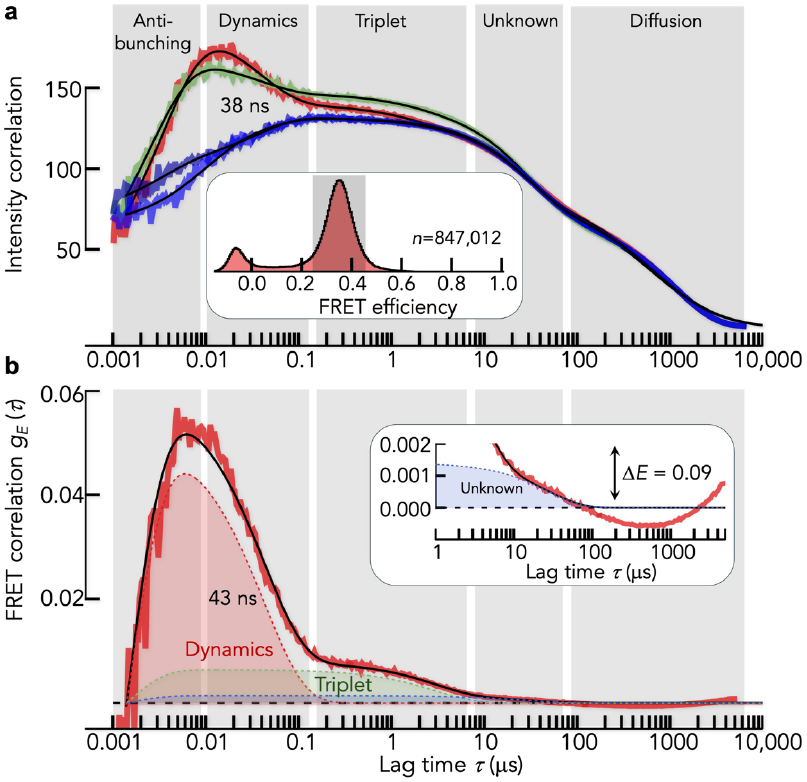
Dynamics of the IDP ΔMyc. **(a)** Acceptor (red) and donor (green) autocorrelation functions and cross-correlation functions (blue) for ΔMyc. The FRET efficiency histogram together with the range of bursts chosen for the calculation of the correlation functions is shown as inset. Vertical gray bars indicate the decay components. Black lines are global fits with 5 decay components (see Methods). **(b)** FRET correlation function computed for the data in (a). Black line is a fit with 4 decays (see Methods). Inset: Zoom at the long-time microsecond regime.

We can translate this amplitude into a lower detection limit for FRET changes. Assuming a 2-state model, the apparent FRET efficiencies ***E***_**1**_ and ***E***_**2**_ of the two conformational states should differ by at least 0.09 FRET efficiency units to overcome the residual diffusion artifacts in FRET correlation functions, i.e., to exceed an amplitude of 0.002.

## Discussion

SmFRET experiments have become an integral part of a ‘new’ era in structural biology that aims at establishing a holistic picture of proteins as dynamic structures^60,61^. The theoretical foundations to rigorously analyze smFRET experiments have been laid over the past two decades^15,41,62-69^. Here, we presented a helpful addition. We showed how the conformational dynamics of single molecules can be retrieved from FRET correlation functions in diffusion-based smFRET experiments. The advantages of the method are fourfold. First, it is simple and can be applied to any smFRET experiment with single-photon detection, irrespective of the data acquisition mode (pulsed- or continuous wave excitation, 2 channel or 4 channel detection). Second, the method effectively suppresses the diffusion component in standard correlation functions, thus providing direct insights into dynamics at timescales from micro-to milliseconds. Third, a few thousand molecules are sufficient to obtain a reasonable signal-to-noise ratio in FRET correlation functions, which substantially reduces measurement times compared to other model-free approaches such as RASP^30^, 2D-FLCS^27,28^, and lifetime-filtered FCS (fFCS)^24,25^. Fourth, the method provides diagnostic tools to identify static heterogeneity, artifacts such as photo-bleaching, and it serves as a rigorous test for model-based fitting approaches that are standardly used in diffusion-based smFRET and integrative structural biology approaches nowadays^9,10,15,19,20,61,70,71^. Hidden-Markov-model fitting of smFRET data with diffusing molecules suffers from our incomplete knowledge on the impact of diffusion through the in-homogeneously illuminated confocal volume and photophysical non-idealities such as triplet dynamics on the photon traces. Are dynamics at tens or hundreds of microseconds poised by the fluctuating photon rate due to triplet and diffusion? Our results demonstrate that albeit these non-idealities can never completely be eradicated, they are substantially suppressed on the FRET efficiency coordinate, which implicitly supports the validity of Markov-model fitting approaches in diffusion-based smFRET experiments. Most importantly, FRET correlation functions provide a model-independent tool to extract dynamic timescales from the photon traces of individual molecules. This is of particular importance for benchmarking Hidden-Markov models. Our tests of the method explored the impact of protein concentrations, bleaching, quenching, triplet, photo-physical saturation, and different detection volumes.

In conclusion, two processes impact FRET correlation functions most: (i) bleaching, which introduces slow decays in the correlation functions, and (ii) triplet dynamics, which causes fast decays at timescales 1 - 10 μs for the Alexa dye-pair used here. With the calculation of cross-correlation ratios we provide the diagnostic tool to identify bleaching. Whereas experiments performed with pulsed interleaved excitation (PIE, ALEX)^45-47^ are ideal to remove bleached molecules, the fluctuations due to triplet transitions are unavoidable. We hope that the simple model-free FRET correlation approach presented here will be a useful addition to the current toolkit of smFRET experiments.

## Methods

### Data simulations and numerical calculations

#### Simulation of Brownian diffusion

To test the performance of our method, we performed simulations of the photon time traces of diffusing molecules that undergo structural dynamics while diffusing through a confocal spot using the Fretica package (https://schuler.bioc.uzh.ch/programs/), developed by Daniel Nettels and Benjamin Schuler (University of Zurich). We obtained trajectories of diffusing particles with Brownian dynamics in spherical coordinates, assuming a radial symmetric confocal volume, which is located at the origin of the coordinate system. The time evolution of particle radial coordinates ***r***(***t***) was obtained with

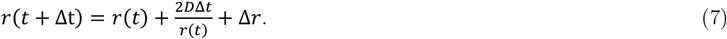

where **Δ*t*** is a timestep, ***D***is the diffusion coefficient, and **Δ*r*** is a stochastic move drawn from a normal distribution with zero mean and variance 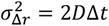. We used **Δ*t* = 1 μs**, except for simulations with the coarse-grained photophysical model (Fig. 5a) for which we used **Δ*t* = 0.5 μs**. The diffusion coefficient of the particles was ***D*= 5 10**^**−5**^ **μm**^**2**^**⁄μs**, which corresponds to a medium-sized protein (Stokes radius of 4.3 nm at room temperature in water). The simulations were initialized by randomly placing particles in a simulation sphere with a radius of ***R* = 3 μm**, afterwards each particle was simulated until it leaves this sphere. The number of initial particles was drawn from a Poisson distribution with a mean ***n***_**0**_ **= 4*πR***^**3**^***c***_**0**_**⁄3**, where ***c***_**0**_ is the bulk particle concentration. The loss of particles that diffuse out of the simulation sphere was compensated by periodically (with period ***T***_***new***_ **= 1000Δ*t***) placing new particles inside the sphere. The number of inserted particles and the distribution of their positions ***c***_***new***_(***r***) were chosen based on the solution of the diffusion equation that describes how an empty sphere fills over time ***T***_***new***_ due to the constant bulk concentration at its outer border

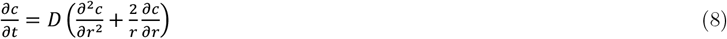

with the initial conditions ***c***(***r* < *R, t* = 0**) **= 0** and the boundary conditions ***c***(***r* = *R, t***) **= *c***_**0**_ and ***c***(***r* → 0, *t***) **= 0**. The solution of eq. 8 is known^72^ and is given by

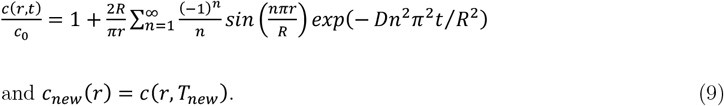

The mean number of particles that enter the sphere ***n***_***new***_ was calculated by integrating ***c***_***new***_(***r***) over the volume of the sphere. After each time interval ***T***_***new***_, a random number of new particles was drawn from the Poisson distribution with mean ***n***_***new***_ and the positions inside the sphere were randomly chosen from the distribution with the density function ***P***_***new***_(***r***) **= 4*πr***^**2**^ ***c***_***new***_(***r***)**⁄*n***_***new***_. In total, we simulated particle trajectories for **1800 s** in most cases whereas **3600 s** long trajectories were used for simulations of a triplet kinetics model.

#### Simulation of conformational dynamics and photon traces

Once the particle trajectories were simulated, we added stochastic conformational dynamics simulated according to the rate equation

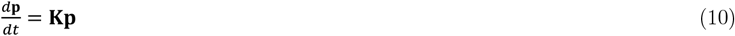

where **p** is the population vector of states and **k** is a matrix that contains the transition rates between states (see below). Each state is characterized by the rate of acceptor and donor photon emission, which is proportional to the intensity profile in the confocal spot

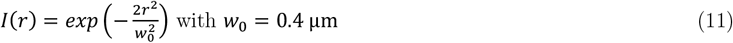

such that the total photon rate at the center of the excitation volume is ***λ***_***tot***_ **= 0.4 μs**^**−1**^. Given the trajectories of positions and conformational states, we simulated the emission of photons and constructed TTTR (time-tagged time-resolved) files. Unless stated otherwise, we used a realistic background photon rate of ***λ***_***d***_ **= 2 10**^**−3**^ **μs**^**−1**^ for the donor channel and ***λ***_***a***_ **= 10**^**−3**^ **μs**^**−1**^ for the acceptor channel. Except for the simulation of non-Markov dynamics (Fig. 3) and in simulations of the coarse-grained photophysical model (Fig. 5a), we introduced different detection efficiencies for the dyes (***γ* = *Q***_***a***_**ζ**_***a***_**⁄*Q***_***d***_**ζ**_***d***_ **= 1.15**), where ***Q***_***a***,***d***_ and **ζ**_***a***,***d***_ are the quantum yields and detection efficiencies for acceptor and donor dye, respectively. Crosstalk (leakage) of donor photons into the acceptor channel (***β***_***DA***_ **= 0.05**) and of acceptor photons into the donor channel (***β***_***AD***_**= 0.003**) together with the probability to directly excite the acceptor with the donor excitation laser (***α* = 0.048**) were also included.

#### Two state model

In our simulations of the 2-state model (Fig. 2, 4a-c), we simulated particles that switch between a low FRET-state 1 and a high FRET-state 2. The two states are characterized by the corrected FRET efficiencies ***ϵ***_**1**_ **= 0.1** and ***ϵ***_**2**_ **= 0.9**. The kinetic rate matrix for this model is

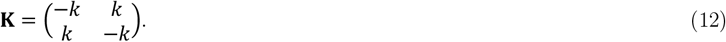

For simplicity, forward and backward rates were equal ***k***. We simulated the systems with wide range of values for ***k*** (0.1, 0.08,0.05, 0.03, 0.025, 0.015, 0.01, 0.008, 0.005, 0.003, 0.002, 0.001, 0.0008, 0.0005, 0.0003, 0.0002, 0.00015, 0.0001, 0.00005) given in units of **μs**^**−1**^. We also simulated the system with ***k* = 0 *μs***^**−1**^ to access static heterogeneity. The initial state for each particle was chosen randomly with equal probabilities to be in low or high FRET state. All the above simulation were done for particle trajectories obtained with different bulk concentrations (25.0, 50.0, 100.0, 200.0, 300.0, 400.0, 500.0), given in units of **pM**. A threshold of 100 photons was used to identify bursts.

#### Non-Markov model and likelihood maximization

Simulations of the photon traces for the 2-state Markov and 4-state non-Markov model were performed as described above. We simulated ‘pure’ systems in the absence of background and instrumental imperfections (***γ* = 1, *β***_***DA***_ **= *β***_***AD***_**= 0, *α* = 0**) such that ***E* = *ϵ*** and identified bursts using a threshold of 50 photons. The rate matrix of the 2-state system is given by eq. 12 and the rate matrix of the 4-state non-Markov system is given by

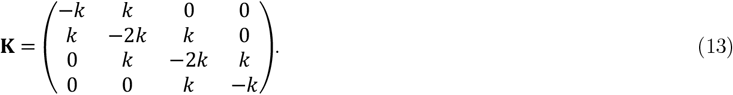

The fit of the simulated photon traces with the 2-state (eq. 12) and 4-state (eq. 13) models was done by maximizing the log-likelihood function of all photon trajectories simultaneously. The log-likelihood function for the ***i*’**s burst with ***N*** photons is given by

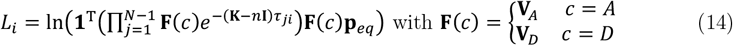

Here, **p**_***eq***_ is the vector of equilibrium probabilities that is the solution of eq. 10 with ***d*p⁄*dt* = 0** where **0** and **1** are vectors of zeros and ones, respectively, ***τ***_***ji***_ is the time between photon ***j*** and ***j* − 1** in the ***i*’**s burst, **I** is the unit matrix, **V**_***A***_ and **V**_***D***_are the detection matrices for acceptor and donor, respectively. For the 2-state model, the detection matrices are

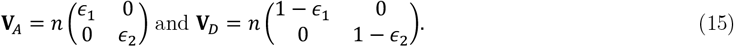

Here, ***n*** is the total photon emission rate of donor and acceptor. We maximized the summed log-likelihood of all ***M*** photon traces given by 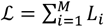 using the Fretica software package. The fitting parameters were ***k, ϵ***_**1**_, ***ϵ***_**2**_, and ***n***. To compute the FRET correlation functions, we re-colored the data by simulating state and colors of the measured photon trajectories with the parameters obtained from the fits and then computed the FRET efficiency histogram and correlation function using the recolored data. To minimize the noise in the recolored FRET correlation functions, we recolored the data ten times and computed the average FRET correlation function of the ten recolored data sets. We also computed the theoretical FRET correlation function of the 4-state model using

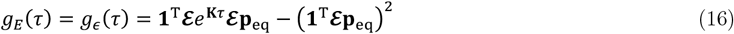

With

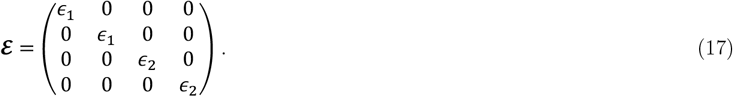

The function is shown as dashed line in Fig. 3d. Notably, eq. 16 is only correct under ideal conditions (no differences in dye brightness, see Supplementary Information), which are fulfilled in this simulation.

#### Simulations with bleaching

To access the effect of bleaching (Fig. 4a-c), we modelled a system that switches between 4 states: donor-only (***D***), acceptor-only (***A***), low FRET (***DA***_**1**_) with ***ϵ***_**1**_ **= 0.1**, and high FRET (***DA***_**2**_) with ***ϵ***_**2**_ **= 0.9**. The rate matrix **k** in this case is a combination of the rate matrix **k**_**0**_ for conformational transitions between ***DA***_**1**_ and ***DA***_**2**_ and the rate matrix **k**_**1**_ that describes photobleaching. Since photobleaching is a function of the excitation intensity, the total rate matrix involves the excitation profile (eq. 11) and is now given by

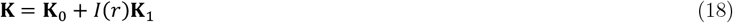

With the matrices

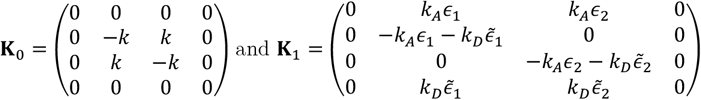

Where 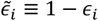. Here, ***k***_***A***_ and ***k***_***D***_are acceptor and donor bleaching rates at the center of the confocal volume (***r* = 0**), respectively. We used realistic bleaching rates of ***k***_***A***_ **= *k***_***D***_**= 8 × 10**^**−4**^ **μs**^**−1**^. The conformational rates were ***k* = 3 × 10**^**−3**^ **μs**^**−1**^. The initial state was chosen randomly according to the probability vector **p**_**0**_ **=** (**0.1 0.4 0.4 0.1**)^**T**^ in the basis **{*D, DA***_**1**_, ***DA***_**2**_, ***A*}**. We also simulated photon emission using a pulsed-interleaved excitation (PIE) scheme^45-47^ of both dyes with ***γ***_***PIE***_ **= 2** to allow the filtering of bleached molecules in the analysis. To this end, we simulated photon emissions after both donor- and acceptor excitation and experimental instrumental response functions (IRF) were used to get realistic arrival time distributions within each PIE period. A threshold of 75 photons was used to identify bursts.

#### Simulation of the coarse-grained photophysical scheme

To understand how the photophysics of real dye pairs impact the photon trajectories of molecules that diffuse through an inhomogeneously illuminated confocal volume, we developed the coarse-grained (CG) model of the kinetic scheme (Fig. 5a) published by Nettels *et. al*. (Supplementary Information).^50^ The original model includes ground singlet states (***S***_**0**_), excited singlet states (***S***_**1**_) and triplet states (***T***_**1**_) for both dyes, resulting in the 9 possible states of the system: **{*S***_**0**_***S***_**0**_, ***S***_**1**_***S***_**0**_, ***S***_**0**_***S***_**1**_, ***S***_**1**_***S***_**1**_, ***T***_**1**_***S***_**0**_, ***T***_**1**_***S***_**1**_, ***S***_**0**_***T***_**1**_, ***S***_**1**_***T***_**1**_, ***T***_**1**_***T***_**1**_**}**. The first and second letter denote the state of donor and acceptor, respectively. Upon absorption of a photon, the dyes undergo transitions form ground to excited singlet states (***S***_**0**_ **→ *S***_**1**_) with the rates ***k***_***ex***_ and ***αk***_***ex***_ for donor and acceptor, respectively, where ***α*** is the probability to directly excite the acceptor at the excitation wavelength of the donor. The value of ***k***_***ex***_ is proportional to the intensity of the incident light and fluctuates due to the diffusion of the particle though the confocal volume. In addition to the classical relaxation pathways ***S***_**1**_ **→ *S***_**0**_, non-radiative interstate crossings ***S***_**1**_ **→ *T***_**1**_ and ***T***_**1**_ **→ *S***_**0**_ and singlet-singlet and singlet-triplet annihilation routes must be considered (Fig. 5a). The timescale of radiative transitions is known to be nanoseconds^50^ whereas intersystem crossings occur at microsecond timescales. In addition, fluctuations of ***k***_***ex***_ due to diffusion are in the order of tens to hundreds of microseconds. Considering this separation of timescales, we built the CG model by assembling states into four groups: ***SS* = {*S***_**0**_***S***_**0**_, ***S***_**1**_***S***_**0**_, ***S***_**0**_***S***_**1**_, ***S***_**1**_***S***_**1**_**}, *TS* = {*T***_**1**_***S***_**0**_, ***T***_**1**_***S***_**1**_**}, *ST*= {*S***_**1**_***T***_**0**_, ***S***_**1**_***T***_**1**_**}**, and ***TT*= {*T***_**1**_***T***_**1**_**}** (Fig. 5a, Supplementary Information). Notably, transitions between these four groups (CG states) corresponds to non-radiative intersystem crossings in the original model. Radiative nanosecond transitions are modelled in form of donor and acceptor photon rates for each CG state, leading to the emission rates 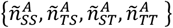 and 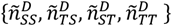 with 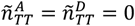. Transition rates between CG-states were obtained by equating the steady-state fluxes between CG states with the sum of fluxes of all corresponding transitions of the full-model (Supplementary Fig. 2). In the realistic range ***k***_***ex***_ **= 0.0 − 0.08 *ns***^**−1**^, we found an approximately linear dependence of the emission rates on ***k***_***ex***_ (Supplementary Fig. 3). Generally, one would expect that also the transition rates of the CG model depend on ***k***_***ex***_. We found that ***S* → *T***transitions, such as ***SS* → *TS*** or ***ST*→ *TT***, depend nearly linearly on ***k***_***ex***_, whereas ***T*→ *S*** transitions, such as ***TS* → *SS*** or ***TT*→ *TS***, are insensitive to changes in ***k***_***ex***_ (Supplementary Fig. 2). This allowed us to write the kinetic rates matrix of the CG model in form of eq. 18. Here, 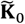contained all ***T*→ *S*** transition rates whereas 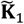 contained the ***S* → *T***transitions that were linearly dependent on ***k***_***ex***_. The dependence of the excitation rate on the location in the confocal volume is given by eq. 11 with ***k***_***ex***_ **= 0.08 *ns***^**−1**^ in the center of simulated confocal spot (***r* = 0, *I***(**0**) **= 1**). The initial state of each simulated particle was ***SS*** as it is the equilibrium state at zero intensity outside the confocal spot. The photon rates of the CG states were obtained from a fit of the CG model to numerical results of the full model and were uniformly adjusted such that the maximum photon rate in the center of the confocal volume is ***λ***_***tot***_ **= 0.4 μs**^**−1**^, which results in a realistic detection rate of 160 photons per millisecond. All parameters of the full and CG model are reported in Supplementary Tables 1-2. A direct comparison between the 9-state model and the CG model is shown in Supplementary Fig. 4.

#### Calculations of FRET correlation functions with different brightness

For the calculation of eq. 5-6 and the numerical results shown in Fig. 4d-j, we used the framework developed by Gopich & Szabo^16^. For a 2-state model, the detection matrices for acceptor and donor are defined by

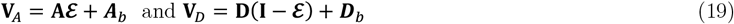

with the matrices

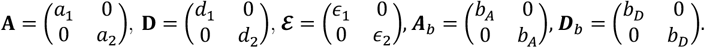

Here, ***ε*** is the matrix of true FRET efficiencies, **A** and **D** are the brightness matrices, containing the brightness values of the states, and ***A***_***b***_ and ***D***_***b***_ contain the background photon counts in the acceptor ***b***_***A***_ and donor ***b***_***D***_ channel. We use eq. 2 to compute the FRET correlation function in this case. To this end, we first compute the photon-pair correlation functions between photons of type X and Y according to

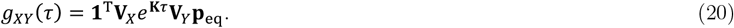

Here, **k** is the rate matrix of the 2-state model (eq. 12) and **p**_**eq**_ is the equilibrium vector of states. The correlation function of all pairs irrespective of color is

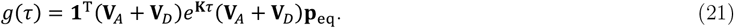

Since ***g***_***XY***_(***τ***) **∝ *N***_***XY***_(***τ***) and ***g***(***τ***) **∝ *N***(***τ***) with 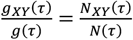, we can use eq. 2 together with the definition of the uncorrected mean FRET efficiency

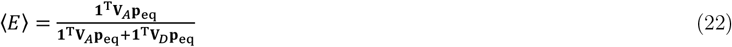

to compute ***g***_***E***_(***τ***). For the special case of state-independent brightness, we set ***a***_**1**_ **= *a***_**2**_ **= *a*** and ***d***_**1**_ **= *d***_**2**_ **= *d*** in eq. 19.

#### Fitting of correlation functions

The experimental FRET correlation functions were fitted with a sum of exponentials with a manually determined offset. The offset was determined based on the final 100 μs of the correlation function. The correlation functions of the DNA samples at pH 4 and pH 7 (Fig. 7b) were fitted globally at each pH value. The global parameters for each pH were the relaxation times and the amplitudes were local parameters. FRET correlation functions from Brownian dynamics simulations with Poisson photon emission statistics (Fig. 2c,e and Fig. 3b,d) were fitted with single exponential function without offset. Given that these decays only show ***τ*** a single decay component, we fitted the data including relative weights for each lag time that were given by the number of total photon pairs at this lag time.

The intensity correlation functions of the nsFCS experiments of ΔMyc (Fig. 8a) were globally fitted with

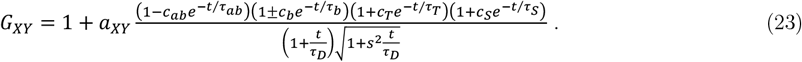

Here, ***X*** and ***Y*** indicate the photon pairs (***AA, DD, AD, DA***), ***a***_***XY***_ is the inverse average number of molecules in the confocal volume, and the indices ***ab, b, T, S*** indicate the amplitudes ***c*** and relaxation times ***τ*** of antibunching, bunching (conformational dynamics), triplet, and unassigned process, respectively. The denominator describes the diffusion were ***τ***_***D***_ is the diffusion time and ***s*** describes the aspect ratio of the confocal volume. Plus and minus in the second factor in the nominator indicate autocorrelation functions (**+**) and cross-correlation functions (**−**).

The FRET correlation function of the nsFCS experiment was fitted with the function

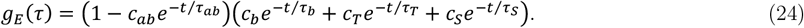

Here, subscripts have the same meaning as in eq. 23.

### Microscope and experimental analysis

#### Single-molecule experiments

The smFRET experiments were performed using a MicroTime 200 (PicoQuant) equipped with an Olympus IX73 inverted microscope. Linearly polarized light from a 485 nm diode laser (LDH-D-C-485, PicoQuant) and unpolarized light at 594 nm from a supercontinuum light source (Solea, PicoQuant) were used to excite donor and acceptor alternatingly with a repetition rate of 20 MHz in an alternating manner. For nsFCS experiments (Fig. 8), we used a continuous excitation of the donor only. In both cases, the excitation light was guided through a major dichroic mirror (ZT 470-491/594 rpc, Chroma) to a 60x, 1.2NA water objective (Olympus) that focused the beam into the sample. A home-made sample cuvette was used with a volume of 50 μl made of round quartz cover slips of 25 mm diameter (Esco Optics) and borosilicate glass 6 mm diameter cloning cylinder (Hilgenberg) using Norland 61 optical adhesive (Thorlabs). We performed the measurements at a laser power of 100 μW (485 nm) and 20 μW (594 nm) measured at the back aperture of the objective. The photons emitted from the sample passed through the same objective and after passing the major dichroic mirror (ZT 470-491/594 rpc, Chroma), the residual excitation light was filtered by a long-pass filter (BLP01-488R, Semrock). The light was then focused on a 100 µm pinhole. The sample fluorescence was detected either with two channels (Fig. 6-7) or four channels (Fig. 8). Donor and acceptor fluorescence was separated via a dichroic mirror (T585 LPXR, Chroma) and each color was focused onto a single-photon avalanche diode (SPAD) (Excelitas) with additional bandpass filters: FF03-525/50, (Semrock) for the donor SPAD and FF02-650/100 (Semrock) for the acceptor SPAD. The arrival time of every detected photon was recorded with a HydraHarp 400M time-correlated single photon counting module (PicoQuant) at a resolution of 32 ps. All measurements were performed at 23°C.

SmFRET experiments on DtpA were measured in 20 mM sodium phosphate pH 7.5, 150 mM NaCl, 0.002% LMNG (lauryl-maltose-neopentyl-glycol), and 20 mM di-thio-threitol (DTT). The experiments on dsDNA were measured in 20 mM sodium phosphate pH 7 or 4, 100 mM β-mercapto-ethanol, 0.001% Tween-20. Experiments with ΔMyc were performed in 20 mM TrisHCl pH 8, 2 M GdmCl, 100 mM β-mercapto-ethanol, and 0.001% Tween-20. Bursts were identified according to standard routines^40,54^ using a photon threshold of 50 (DtpA), 50 (dsDNA), and 150 (ΔMyc).

#### Burst selection and bleaching filter

The raw photon numbers for donor (***n*′**_***D***_) and acceptor (***n*′**_***D***_) were corrected for background, quantum yields and detection efficiencies of the microscope (***γ* = 1.12 ± 0.09**), cross-talk (***β***_***DA***_ **= 0.050 ± 0.003** and ***β***_***AD***_**= 0.0021 ± 0.0004**), and acceptor direct excitation (***α* = 0.049**), leading to the corrected photon numbers ***n***_***DD***_and ***n***_***DA***_, where the first subscript indicates the excitation and the second indicates emission. The corrected mean FRET efficiency (temporal average over the burst duration) of a burst is then

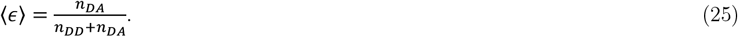

For DtpA, dsDNA, and the results of the simulations including bleaching (Fig. 4a-c, right), bursts containing no acceptor fluorescence after acceptor direct excitation (donor-only) were removed by only selecting bursts with a dye stoichiometry ratio **0.25 ≤ *S* ≤ 0.75** where the stoichiometry of a burst is defined by

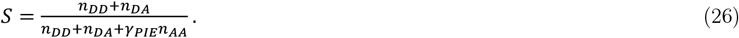

Here, ***γ***_***PIE***_ **= 2 − 2.5** is a factor that accounts for the different excitation intensities for donor and acceptor. The remaining bursts were additionally filtered to exclude bursts in which the acceptor bleached during the transit through the confocal volume. To this end, we only selected those bursts for which the mean arrival times after donor **⟨*t***_***Dex***_**⟩** and acceptor **⟨*t***_***Aex***_**⟩** excitation were similar. We then define the burst asymmetry as

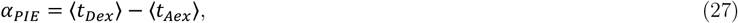

We chose a very restrictive threshold of ***α***_***PIE***_ **≤ 0.05 − 0.1 *ms*** to exclude bleaching artifacts in the FRET correlation functions.

