## Supplementary Information for "Model-free photon analysis of diffusion-based single-molecule FRET experiments"

**1.1 Derivation of the FRET correlation function $g_E(\tau)$: General case.** The structural dynamics of individual biomolecules results in stochastic fluctuations of the distance $r(t)$ between the donor and acceptor dyes in time $t$. We assume that $r(t)$ is a stationary process. Within the framework of Förster theory, this distance is related to the theoretical 'true' FRET efficiency $\varepsilon(t)$ via

$$\varepsilon(t) = \frac{R_0^6}{R_0^6 + r(t)^6}, \tag{S1}$$

which is a quantity that changes continously in time. Here, $R_0$ is the dye-specific Förster distance (5.4 nm for the dye pair AlexaFluor 488 and AlexaFluor 594 used in this study)[1]. Clearly, $\varepsilon(t)$ is also a stochastic variable and the natural approach to characterize the fluctuations in $\varepsilon(t)$ is based on the autocorrelation function

$$g_\varepsilon(\tau) = \langle[\varepsilon(0) - \langle\varepsilon\rangle][\varepsilon(\tau) - \langle\varepsilon\rangle]\rangle_\tau = \langle\varepsilon(0)\varepsilon(\tau)\rangle_\tau - \langle\varepsilon\rangle^2. \tag{S2}$$

Here, $\tau$ is the lag time and the brackets $\langle\dots\rangle_\tau$ indicate averaging over all pairs of $\varepsilon$ values that are separated by $\tau$. The bracket $\langle\dots\rangle$ indicates the average over the whole ensemble of possible values of $\varepsilon$. We define the ensemble average $\langle\varepsilon\rangle$ as

$$\langle\varepsilon\rangle = \int_0^1 d\varepsilon\, f(\varepsilon)\varepsilon \text{ with the normalization } \int_0^1 d\varepsilon\, f(\varepsilon) = 1. \tag{S3}$$

Here, $f(\varepsilon)$ is the distribution of $\varepsilon$ over the ensemble. To define the correlation function $\langle\varepsilon(0)\varepsilon(\tau)\rangle_\tau$, we introduce the joint distribution $F(\varepsilon_1, \varepsilon_2|\tau)$ for two random variables $\varepsilon_1 = \varepsilon(0)$ and $\varepsilon_2 = \varepsilon(\tau)$ that are separated by the time $\tau$. With this definition, the autocorrelation function is given by

$$\langle\varepsilon(0)\varepsilon(\tau)\rangle_\tau = \int_0^1\int_0^1 d\varepsilon_1 d\varepsilon_2 F(\varepsilon_1, \varepsilon_2|\tau)\varepsilon_1\varepsilon_2 \text{ with } \int_0^1\int_0^1 d\varepsilon_1 d\varepsilon_2 F(\varepsilon_1, \varepsilon_2|\tau) = 1. \quad \text{(S4)}$$

In an experiment however, we neither observe $\varepsilon(t)$ nor can we compute the correlation function given by eq. S4. Instead, in smFRET, we detect photons from the donor and acceptor. For each acceptor photon, we assign an apparent FRET efficiency $E = 1$ and each donor photon obtains $E = 0$. Each burst is then a stochastic trajectory of 0's and 1's, which is a realization of the process $E(t)$. To derive the correlation function of this process, we first define the mean value of $E$. In the ideal case, i.e., no background, identical brightness of both dyes, no triplet-state dynamics and antibunching, and in the absence of spectral crosstalk and any microscope imperfection, we can define the probability that a detected photon is of type $E$ given that the true FRET efficiency is $\varepsilon$. This probability is given by

$$p(E|\varepsilon) = \begin{cases} p(1|\varepsilon) = \varepsilon \\ p(0|\varepsilon) = 1 - \varepsilon \end{cases} \text{ such that } \sum_{E=0}^{1} p(E|\varepsilon) = 1. \quad \text{(S5)}$$

In real experiments however, we have to take the non-idealities into account. To this end, we introduce the detection functions $\Omega_A(\varepsilon)$ and $\Omega_D(\varepsilon)$ for the acceptor and donor channel, which turns the probabilities in eq. S5 into weights

$$w(E|\varepsilon) = \begin{cases} w(1|\varepsilon) = \varepsilon \cdot \Omega_A(\varepsilon) \\ w(0|\varepsilon) = (1 - \varepsilon) \cdot \Omega_D(\varepsilon) \end{cases}. \quad \text{(S6)}$$

Note that these weights do not necessarily sum to one. With these weights, the measured mean FRET efficiency is given by

$$\langle E\rangle = \frac{1}{S_E}\int_0^1 d\varepsilon f(\varepsilon) \sum_{E=0}^{1} w(E|\varepsilon)\, E = \frac{1}{S_E}\int_0^1 d\varepsilon f(\varepsilon)\, \varepsilon \cdot \Omega_A(\varepsilon) \quad \text{(S7)}$$

with the normalization

$$S_E = \int_0^1 d\varepsilon f(\varepsilon) \sum_{E=0}^{1} w(E|\varepsilon) = \int_0^1 d\varepsilon f(\varepsilon) \left[\Omega_D(\varepsilon) + \varepsilon\left(\Omega_A(\varepsilon) - \Omega_D(\varepsilon)\right)\right]. \quad \text{(S8)}$$

To define the autocorrelation function for the measured FRET efficiency, we analogously introduce weights for photon pairs. The weights $W(E_1, E_2|\varepsilon_1, \varepsilon_2, \tau)$ for the case of two detected photons of type $E_1$ and $E_2$ given the lag time $\tau$ and the corresponding true FRET efficiencies $\varepsilon_1$ and $\varepsilon_2$, are given by

$$W(1,1|\varepsilon_1,\varepsilon_2,\tau) \;=\; \varepsilon_1\varepsilon_2 \cdot \Omega_{AA}(\varepsilon_1,\varepsilon_2,\tau)$$

$$W(0,0|\varepsilon_1,\varepsilon_2,\tau) \;=\; (1-\varepsilon_1)(1-\varepsilon_2) \cdot \Omega_{DD}(\varepsilon_1,\varepsilon_2,\tau)$$

$$W(1,0|\varepsilon_1,\varepsilon_2,\tau) \;=\; \varepsilon_1(1-\varepsilon_2) \cdot \Omega_{AD}(\varepsilon_1,\varepsilon_2,\tau)$$

$$W(0,1|\varepsilon_1,\varepsilon_2,\tau) \;=\; (1-\varepsilon_1)\varepsilon_2 \cdot \Omega_{DA}(\varepsilon_1,\varepsilon_2,\tau). \qquad \text{(S9)}$$

Here, $\Omega_{XY}(\varepsilon_1,\varepsilon_2,\tau)$ are the detection functions for the photon pairs $\{X,Y\}$. In the ideal case (defined above), we would have $\Omega_{XY}(\varepsilon_1,\varepsilon_2,\tau)=1$. The detection functions account for all non-idealities and might even be time-correlated such as in the presence of triplet dynamics or antibunching. With these weights, the autocorrelation function, which we call the FRET correlation function in the main text, is given by

$$g_E(\tau) = \langle [E(0)-\langle E\rangle][E(\tau)-\langle E\rangle]\rangle_\tau =$$
$$\frac{1}{S_{EE}(\tau)}\int_0^1\int_0^1 d\varepsilon_1 d\varepsilon_2 F(\varepsilon_1,\varepsilon_2|\tau)\sum_{E_1=0}^{1}\sum_{E_2=0}^{1} W(E_1,E_2|\varepsilon_1,\varepsilon_2,\tau)(E_1-\langle E\rangle)(E_2-\langle E\rangle) \qquad \text{(S10)}$$

with the normalization

$$S_{EE}(\tau) = \int_0^1\int_0^1 d\varepsilon_1 d\varepsilon_2 F(\varepsilon_1,\varepsilon_2|\tau)\sum_{E_1=0}^{1}\sum_{E_2=0}^{1} W(E_1,E_2|\varepsilon_1,\varepsilon_2,\tau)\;. \qquad \text{(S11)}$$

Here again, $\tau$ is the lag time and the brackets $\langle\ldots\rangle_\tau$ now indicate averaging over all pairs of detected photons of value $E$ that are separated by $\tau$. After expanding the brackets in eq. S10, we write the autocorrelation function as

$$g_E(\tau) = (1-\langle E\rangle)^2\Gamma_{AA}(\tau) + \langle E\rangle^2\Gamma_{DD}(\tau) - \langle E\rangle(1-\langle E\rangle)[\Gamma_{AD}(\tau)+\Gamma_{DA}(\tau)] \qquad \text{(S12)}$$

With eqs. S9, the functions $\Gamma_{XY}(\tau)$ of photon pairs of types $\{X,Y\}$ separated by time $\tau$ are given by

$$\Gamma_{AA}(\tau) = \frac{1}{S_{EE}(\tau)}\int_0^1\int_0^1 d\varepsilon_1 d\varepsilon_2 F(\varepsilon_1,\varepsilon_2|\tau)\varepsilon_1\varepsilon_2\Omega_{AA}(\varepsilon_1,\varepsilon_2,\tau) = \langle E(0)E(\tau)\rangle_\tau$$

$$\Gamma_{DD}(\tau) = \frac{1}{S_{EE}(\tau)}\int_0^1\int_0^1 d\varepsilon_1 d\varepsilon_2 F(\varepsilon_1,\varepsilon_2|\tau)(1-\varepsilon_1)(1-\varepsilon_2)\Omega_{DD}(\varepsilon_1,\varepsilon_2,\tau) = \langle [1-E(0)][1-E(\tau)]\rangle_\tau$$

$$\Gamma_{AD}(\tau) = \frac{1}{S_{EE}(\tau)}\int_0^1\int_0^1 d\varepsilon_1 d\varepsilon_2 F(\varepsilon_1,\varepsilon_2|\tau)\varepsilon_1(1-\varepsilon_2)\Omega_{AD}(\varepsilon_1,\varepsilon_2,\tau) = \langle E(0)[1-E(\tau)]\rangle_\tau$$

$$\Gamma_{DA}(\tau) = \frac{1}{S_{EE}(\tau)}\int_0^1\int_0^1 d\varepsilon_1 d\varepsilon_2 F(\varepsilon_1,\varepsilon_2|\tau)(1-\varepsilon_1)\varepsilon_2\Omega_{AA}(\varepsilon_1,\varepsilon_2,\tau) = \langle [1-E(0)]E(\tau)\rangle_\tau. \qquad \text{(S13)}$$

These terms are nothing else than the fractions of photon pairs of the four possible types, which, for unlimited number of photon pairs, makes eq. 2 in the main text identical to eq. S12. Based on the second equalities in eq. S13, we notice that $\Gamma_{AA}(\tau) + \Gamma_{DD}(\tau) + \Gamma_{AD}(\tau) + \Gamma_{DA}(\tau) = 1$, which is the relation given in eq. 4 in the main text. Importantly, unlike in eq. S2, we would like to note that generally

$$g_E(\tau) = \langle [E(0) - \langle E\rangle][E(\tau) - \langle E\rangle]\rangle_\tau \neq \langle E(0)E(\tau)\rangle_\tau - \langle E\rangle^2. \quad \text{(S14)}$$

Let us investigate the conditions at which an equality in eq. S14 would be justified. We multiply the brackets in eq. S12 and recollect the terms in powers of $\langle E\rangle$, which gives

$$g_E(\tau) = \Gamma_{AA}(\tau) - \big(2\Gamma_{AA}(\tau) + \Gamma_{AD}(\tau) + \Gamma_{DA}(\tau)\big)\langle E\rangle + \big(\Gamma_{AA}(\tau) + \Gamma_{DD}(\tau) + \Gamma_{AD}(\tau) + \Gamma_{DA}(\tau)\big)\langle E\rangle^2. \quad \text{(S15)}$$

Using the second equalities in eq. S13, we find

$$g_E(\tau) = \langle E(0)E(\tau)\rangle_\tau - [\langle E(0)\rangle_\tau + \langle E(\tau)\rangle_\tau]\langle E\rangle + \langle E\rangle^2. \quad \text{(S16)}$$

To arrive at $g_E(\tau) = \langle E(0)E(\tau)\rangle_\tau - \langle E\rangle^2$, one would require that

$$\langle E(0)\rangle_\tau = \langle E(\tau)\rangle_\tau = \langle E\rangle\ , \quad \text{(S17)}$$

which is the hidden assumption in $\langle [E(0) - \langle E\rangle][E(\tau) - \langle E\rangle]\rangle_\tau = \langle E(0)E(\tau)\rangle_\tau - \langle E\rangle^2$. If the identities in eq. S17 were generally true, the correct FRET correlation function could be computed solely from the acceptor autocorrelation ratio and the average FRET efficiency via $g_E(\tau) = \Gamma_{AA}(\tau) - \langle E\rangle^2$ instead of using the more complicated expression in eq. 2 (main text) and S12-13. In a real experiments however, $\langle E(0)\rangle_\tau$, $\langle E(\tau)\rangle_\tau$, and $\langle E\rangle$ differ as soon as the total photon rate is not independent of $\varepsilon(t)$, for instance, if the dye brightness changes with the conformational state of a protein. What is the meaning of $\langle E(0)\rangle_\tau$ and $\langle E(\tau)\rangle_\tau$? For all photon pairs separated by $\tau$, $\langle E(0)\rangle_\tau$ is the mean FRET efficiency computed only from the first photons in the pairs whereas $\langle E(\tau)\rangle_\tau$ is the mean FRET efficiency computed from the second photon in the pairs. Using the second equalities in eqs. S13, we find that

$$\langle E(0)\rangle_\tau = \Gamma_{AA}(\tau) + \Gamma_{AD}(\tau) \text{ and } \langle E(\tau)\rangle_\tau = \Gamma_{AA}(\tau) + \Gamma_{DA}(\tau). \quad \text{(S18)}$$

For a stationary process, we would expect $\Gamma_{AD}(\tau) = \Gamma_{DA}(\tau)$ and therefore $\langle E(0)\rangle_\tau = \langle E(\tau)\rangle_\tau$. However, the averages might depend on the lag time $\tau$, in contrast to the average over all photons $\langle E\rangle$. We demonstrate this aspect with a common case in smFRET, the case in which the brightness of the donor $(d)$ differs from that of the acceptor $(a)$, thus leading to a correction factor $\gamma = a/d \neq 1$. In this case, the weights defined in eq. S9 are given by

$$W(1,1|\varepsilon_1,\varepsilon_2,\tau) \; = \; \varepsilon_1\varepsilon_2 a^2$$

$$W(0,0|\varepsilon_1,\varepsilon_2,\tau) \; = \; (1-\varepsilon_1)(1-\varepsilon_2)d^2$$

$$W(1,0|\varepsilon_1,\varepsilon_2,\tau) \; = \; \varepsilon_1(1-\varepsilon_2)\, ad$$

$$W(0,1|\varepsilon_1,\varepsilon_2,\tau) \; = \; (1-\varepsilon_1)\varepsilon_2 da. \qquad \text{(S19)}$$

Inserting these weights into eq. S13 and using eq. S18, we find

$$\langle E(0)\rangle_\tau = \frac{\int_0^1\int_0^1 d\varepsilon_1 d\varepsilon_2 F(\varepsilon_1,\varepsilon_2|\tau)[\varepsilon_1 ad+\varepsilon_1\varepsilon_2 d(a-d)]}{\int_0^1\int_0^1 d\varepsilon_1 d\varepsilon_2 F(\varepsilon_1,\varepsilon_2|\tau)[d^2+(\varepsilon_1+\varepsilon_2)d(a-d)+\varepsilon_1\varepsilon_2(a-d)^2]}\;. \qquad \text{(S20)}$$

With the definition of the correlation function of the true FRET efficiency (eq. S4), we can rewrite this expression as

$$\langle E(0)\rangle_\tau = \frac{\langle\varepsilon\rangle ad + d(a-d)\langle\varepsilon(0)\varepsilon(\tau)\rangle_\tau}{d^2+2d(a-d)\langle\varepsilon\rangle+(a-d)^2\langle\varepsilon(0)\varepsilon(\tau)\rangle_\tau}\;, \qquad \text{(S21)}$$

which indeed is a lag-time dependent quantity. For completeness, the average FRET eficiency over the whole ensemble of photons is given by

$$\langle E\rangle = \frac{a\langle\varepsilon\rangle}{d+(a-d)\langle\varepsilon\rangle}\;. \qquad \text{(S22)}$$

Hence, already in the very common case of $\gamma \neq 1$, we find $\langle E(0)\rangle_\tau \neq \langle E\rangle$. It is easy to see that for $\gamma = 1$ ($a = d$), $\langle E(0)\rangle_\tau = \langle E\rangle$.[1]

**Derivation of the FRET correlation function $g_E(\tau)$: Ideal case.** Here, we show that that $g_E(\tau) = g_\varepsilon(\tau)$ in the ideal case. The ideal case is given by

$$\Omega_{AA}(\varepsilon_1,\varepsilon_2,\tau) = \Omega_{DD}(\varepsilon_1,\varepsilon_2,\tau) = \Omega_{AD}(\varepsilon_1,\varepsilon_2,\tau) = \Omega_{DA}(\varepsilon_1,\varepsilon_2,\tau) = 1.$$

and

$$\Omega_A(\varepsilon) = \Omega_D(\varepsilon) = 1. \qquad \text{(S23)}$$

For this case, we already demonstrated that

[1] We note that there is an alternative definition of the FRET correlation function as an autocovariance $g_E(\tau) = \langle[E(0)-\langle E(0)\rangle_\tau][E(\tau)-\langle E(\tau)\rangle_\tau]\rangle$, which would lead to $g_E(\tau) = \Gamma_{AA}(\tau)\Gamma_{DD}(\tau) - \Gamma_{AD}(\tau)\Gamma_{DA}(\tau)$ instead of eq. S12. We prefer eq. S12 because $\langle \mathrm{E}\rangle$ is a more natural quantity to define the mean FRET efficiency.

$$g_E(\tau) = \langle E(0)E(\tau)\rangle_\tau - \langle E\rangle^2 = \Gamma_{AA}(\tau) - \langle E\rangle^2 \tag{S24}$$

With the definition of the mean FRET efficiency given by eq. S7-8 and the definition of the mean of the true FRET efficiency in eq. S3 together with our definition of the ideal case (eq. S23), it is obvious that $\langle E\rangle = \langle \varepsilon\rangle$. With eq. S11, S13, and the conditions in S23, we find

$$\Gamma_{AA}(\tau) = \frac{1}{S_{EE}(\tau)}\int_0^1\int_0^1 d\varepsilon_1 d\varepsilon_2 F(\varepsilon_1,\varepsilon_2|\tau)\varepsilon_1\varepsilon_2 \text{ and } S_{EE}(\tau) = 1. \tag{S25}$$

Comparing eq. S25 with eq. S4, also shows that $\langle E(0)E(\tau)\rangle_\tau = \langle \varepsilon(0)\varepsilon(\tau)\rangle_\tau$. Hence, under ideal conditions, we find $g_E(\tau) = g_\varepsilon(\tau)$.

**2. Cancelation of diffusion in FRET correlation functions.** The probability to detect a photon from a molecule depends on the size and shape of the excitation and detection volumes of the microscope. Generally, these volumes are wavelength-dependent and therefore differ between excitation and donor/acceptor emission. To access how these volumes impact regular FCS experiments and particularly the FRET correlation function, we first collect all components that contribute to the photon detection rates of donor and acceptor. Since excitation, Förster-transfer and photon emission are processes that are much faster than the diffusion through the confocal volume, the probability that a molecule at location $\mathbf{r}$ and time $t$ emits a photon can be represented as a product of three factors: (i) the probability of absorbing a photon and exciting the donor, (ii) the probability of emitting either a donor or an acceptor photon, and (iii) the probability of detecting the acceptor or donor photon. The photon rates are therefore given by three factors

$$n_A(\mathbf{r},t) = \underbrace{\sigma I(\mathbf{r})C(\mathbf{r},t)}_{exitation}\quad \underbrace{\varepsilon(t)Q_A(t)}_{transfer+emission}\quad \underbrace{\xi_A T_A(\mathbf{r})}_{detection}$$

$$n_D(\mathbf{r},t) = \underbrace{\sigma I(\mathbf{r})C(\mathbf{r},t)}_{exitation}\quad \underbrace{[1-\varepsilon(t)]Q_D(t)}_{no\ transfer+emission}\quad \underbrace{\xi_D T_D(\mathbf{r})}_{detection}\ . \tag{S26}$$

Here, $\sigma$ is the absorption cross-section of the donor, $I(\mathbf{r})$ is the intensity of the excitation light at position $\mathbf{r}$, $C(\mathbf{r},t)$ is the probability density (concentration) to find a molecule in the volume element $d\mathbf{r}$ around position $\mathbf{r}$ at time $t$, $\varepsilon(t)$ is the true FRET efficiency at time $t$, $Q_A(t)$ and $Q_D(t)$ are the quantum yields of acceptor and donor, respectively. We consider the general case here in which the quantum yields can fluctuate due to quenching processes. Furthermore, $\xi_A$ and $\xi_D$ are the detection efficiencies of the detectors in the acceptor and donor channels, respectively, and $T_A(\mathbf{r})$ and $T_D(\mathbf{r})$ are the profiles of the acceptor and donor detection volumes of the microscope, respectively. It is convenient to separate the internal emission dynamics of a molecule from the terms that depend on the diffusion of the molecule through the confocal volume. We therefore introduce the time-dependent brightnesses of both dyes as

$$a(t) = \xi_A Q_A(t) \text{ and } d(t) = \xi_D Q_D(t) \tag{S27}$$

and we collect the remaining contributions in the terms

$$n_A^o(\mathbf{r}, t) = \sigma I(\mathbf{r}) T_A(\mathbf{r}) C(\mathbf{r}, t) \text{ and } n_D^o(\mathbf{r}, \mathrm{t}) = \sigma I(\mathbf{r}) T_D(\mathbf{r}) C(\mathbf{r}, t)\ . \tag{S28}$$

With this notation, the photon rates can be written as

$$n_A(\mathbf{r}, t) = \varepsilon(t) a(t) n_A^o(\mathbf{r}, t) \text{ and } n_D(\mathbf{r}, t) = [1 - \varepsilon(t)] d(t) n_D^o(\mathbf{r}, t). \tag{S29}$$

In the absence of photo-physical saturation, the brightnesses $a(t)$ and $d(t)$ together with the true FRET efficiency $\varepsilon(t)$ do not depend on the intensity of the excitation light. Hence, $a(t)$, $d(t)$, and $\varepsilon(t)$ are statistically independent of the processes $n_A^o(\mathbf{r}, t)$ and $n_D^o(\mathbf{r}, t)$. This allows us to factorize the intensity auto- and cross-correlation functions of the rates in the following way

$$g_{AA}(\tau) = \langle n_A(\mathbf{r}_1, t)\, n_A(\mathbf{r}_2, t + \tau)\rangle = \langle \varepsilon(t)\, a(t) \varepsilon(t + \tau)\, a(t + \tau)\rangle \cdot \langle n_A^o(\mathbf{r}_1, t) n_A^o(\mathbf{r}_2, t + \tau)\rangle$$

$$g_{DD}(\tau) = \langle n_D(\mathbf{r}_1, t)\, n_D(\mathbf{r}_2, t + \tau)\rangle = \langle [1 - \varepsilon(t)] d(t) [1 - \varepsilon(t + \tau)] d(t + \tau)\rangle \cdot \langle n_D^o(\mathbf{r}_1, t) n_D^o(\mathbf{r}_2, t + \tau)\rangle$$

$$g_{AD}(\tau) = \langle n_A(\mathbf{r}_1, t)\, n_D(\mathbf{r}_2, t + \tau)\rangle = \langle \varepsilon(t) a(t) [1 - \varepsilon(t + \tau)] d(t + \tau)\rangle \cdot \langle n_A^o(\mathbf{r}_1, t) n_D^o(\mathbf{r}_2, t + \tau)\rangle$$

$$g_{DA}(\tau) = \langle n_D(\mathbf{r}_1, t)\, n_A(\mathbf{r}_2, t + \tau)\rangle = \langle [1 - \varepsilon(t)] d(t) \varepsilon(t + \tau) a(t + \tau)\rangle \cdot \langle n_D^o(\mathbf{r}_1, t) n_A^o(\mathbf{r}_2, t + \tau)\rangle\ . \tag{S30}$$

The brackets $\langle \dots \rangle$ in the first factor indicate averaging over internal states of the molecules, whereas as brackets in the second factor indicates averaging of the positions inside the confocal volume ($\mathbf{r}_1$ and $\mathbf{r}_2$). The correlation function of the total intensity is given by

$$g(\tau) = \langle [n_D(\mathbf{r}_1, t) + n_A(\mathbf{r}_1, t)]\, [n_D(\mathbf{r}_2, t + \tau) + n_A(\mathbf{r}_2, t + \tau)]\rangle = g_{AA}(\tau) + g_{DD}(\tau) + g_{AD}(\tau) + g_{DA}(\tau)\ . \tag{S31}$$

From eq. S30, it is easy to see that the terms that contain information on the diffusion through the confocal volume occur as multipliers (second factor). Hence, in the limit in which the detection volumes of donor and acceptor are identical, i.e., $n_D^o(\mathbf{r}, t) = n_A^o(\mathbf{r}, t) \equiv n^o(\mathbf{r}, t)$, the correlation function of the total intensity is

$$g(\tau) = \langle q(t) q(t + \tau)\rangle \langle n^o(\mathbf{r}_1, t) n^o(\mathbf{r}_2, t)\ \rangle$$

with

$$q(t) = d(t) + [a(t) - d(t)] \varepsilon(t). \tag{S32}$$

In the ratios in eq. 3 (main text), the factor $\langle n^o(\mathbf{r}_1,t)n^o(\mathbf{r}_2,t)\,\rangle$ will therefore cancel, thus removing the diffusion term entirely. In reality however, the limit $n_D^o(\mathbf{r},t) = n_A^o(\mathbf{r},t)$ is not fullfilled. The detection volumes $T_A(\mathbf{r})$ and $T_D(\mathbf{r})$ in eq. S25 depend on the detection wavelength, which typically differs by ~100 nm between donor and acceptor dyes in smFRET experiments. Yet, this difference is less problematic if we realize that the product between excitation and detection volume is the relevant quantity, namely $I(\mathbf{r})T_A(\mathbf{r})$ and $I(\mathbf{r})T_D(\mathbf{r})$. In words: either dye can only emit photons if it has been excited first. The product with the excitation volume $I(\mathbf{r})$ therefore introduces a filter that diminishes the difference between $n_D^o(\mathbf{r},t)$ and $n_A^o(\mathbf{r},t)$. To demonstrate this filtering, we use the common Gaussian approximation for excitation and detection volumes. With $\mathbf{r} = (x \quad y \quad z)$, we can write

$$I(\mathbf{r}) \propto \exp\left[-\frac{2(x^2+y^2)}{\omega_{ex}^2}\right]\exp\left[-\frac{2z^2}{z_{ex}^2}\right] \qquad \text{(excitation profile)}$$

$$T_A(\mathbf{r}) \propto \exp\left[-\frac{2(x^2+y^2)}{\omega_A^2}\right]\exp\left[-\frac{2z^2}{z_A^2}\right] \qquad \text{(acceptor detection profile)}$$

$$T_D(\mathbf{r}) \propto \exp\left[-\frac{2(x^2+y^2)}{\omega_D^2}\right]\exp\left[-\frac{2z^2}{z_D^2}\right] \qquad \text{(donor detection profile).} \qquad \text{(S33)}$$

Here, $\omega_{ex}$ and $z_{ex}$ are the lateral and longitudinal dimensions of the excitation volume, respectively, and ($\omega_A$, $z_A$) and ($\omega_D$, $z_D$) are the corresponding dimensions of the detection volumes for acceptor and donor, respectively. Since the multiplication of two Gaussians is again a Gaussian, the dimensions of the products $I(\mathbf{r})T_A(\mathbf{r})$ and $I(\mathbf{r})T_D(\mathbf{r})$ are, using the subscript $X = \{A, D\}$,

$$\tilde{\omega}_X = \left(\frac{1}{\omega_{ex}^2}+\frac{1}{\omega_X^2}\right)^{-1/2} \text{ and } \tilde{z}_X = \left(\frac{1}{z_{ex}^2}+\frac{1}{z_X^2}\right)^{-1/2}. \qquad \text{(S34)}$$

These parameters then determine the effective detection volume $V_{eff} \propto \tilde{\omega}_X^2\tilde{z}_X$. Since the dimensions in a focussed beam scale linearly with the wavelength of the focussed light, we can compute the ratio of the effective detection volumes of donor and acceptor:

$$\frac{V_A}{V_D} = \frac{\tilde{\omega}_A^2\tilde{z}_A}{\tilde{\omega}_D^2\tilde{z}_D} = \left(\frac{\lambda_{ex}^{-2}+\lambda_D^{-2}}{\lambda_{ex}^{-2}+\lambda_A^{-2}}\right)^{3/2}. \qquad \text{(S35)}$$

A calculation for the wavelengths used for our dye pair ($\lambda_{ex} = 485\,nm$, $\lambda_D = 520\,nm$, $\lambda_A = 620\,nm$) is shown in Supplementary Fig. 1. It shows that while the detection volumes of donor and acceptor differ by more than 50%, the effective detection volumes only differ by 20%. Notably, this difference is likely further reduced by focussing the emission light onto a pinhole, which removes out-of-focus light.

Yet, even a 20% difference in the detection volumes of donor and acceptor causes a FRET change simply due to the diffusion of the molecule through the confocal volume. We therefore checked the impact of this effect on the FRET correlation functions. To exclusively focus on the impact of different detection volumes, we consider the case $\varepsilon(t) = 0.5$, i.e., a molecule with no internal dynamics and with identical brightness of donor and acceptor ($a = d = \gamma = 1$). The correlation functions for the four possible photon pairs are given by eqs. S25. With our assumptions, the first factor in these equations reduces to a constant ($g_0 = 1/4$) whereas the second factor is an average over all spatial locations in the confocal volume, thus leading to

$$g_{XY}(\tau) = g_0 \iint d\mathbf{r}_1 d\mathbf{r}_2\, I(\mathbf{r}_1)T_X(\mathbf{r}_1)\, I(\mathbf{r}_2)T_Y(\mathbf{r}_2)\, \langle C(\mathbf{r}_1,t)C(\mathbf{r}_2,t+\tau)\rangle \tag{S36}$$

where the subscripts $X$ and $Y$ indicate the color of the photon. The term $\langle C(\mathbf{r}_1,t)C(\mathbf{r}_2,t+\tau)\rangle$ is the probability density to find a molecule at $\mathbf{r}_1$ at time $t$ and at $\mathbf{r}_2$ at time $t+\tau$. This probability density consists of two components, (i) the probability of having two distinct molecules in the confocal volume and (ii) the probability that the molecule diffuses from $\mathbf{r}_1$ to $\mathbf{r}_2$ within the time lag $\tau$. With the Green's function of the diffusion equation we have

$$\langle C(\mathbf{r}_1,t)C(\mathbf{r}_2,t+\tau)\rangle = \langle C\rangle^2 + \frac{\langle C\rangle}{(4\pi D\tau)^{3/2}} \exp\left[-\frac{|\mathbf{r}_1-\mathbf{r}_2|^2}{4D\tau}\right]. \tag{S37}$$

Here, $D$ is the translational diffusion coefficient of the molecule and $\langle C\rangle$ is the bulk concentration of molecules. In a single-molecule setup, the chance of finding two molecules simultaneously in the confocal volume is low, which allows us to drop the first term in eq. S37. Then substituting eq. S37 into eq. S36 and using $\mathbf{r}_1 = (x_1 \quad y_1 \quad z_1)$ and $\mathbf{r}_2 = (x_2 \quad y_2 \quad z_2)$, gives

$$g_{XY}(\tau) = \frac{g_0\langle C\rangle}{(4\pi D\tau)^{3/2}} \iint d\mathbf{r}_1 d\mathbf{r}_2 \exp\left[-\frac{2(x_1^2+y_1^2)}{\tilde{\omega}_X^2}\right] \exp\left[-\frac{2z_1^2}{\tilde{z}_X^2}\right] \exp\left[-\frac{2(x_2^2+y_2^2)}{\tilde{\omega}_Y^2}\right] \exp\left[-\frac{2z_2^2}{\tilde{z}_Y^2}\right] \exp\left[-\frac{|\mathbf{r}_1-\mathbf{r}_2|^2}{4D\tau}\right]. \tag{S38}$$

Solving the integrals gives

$$g_{XY}(\tau) = g_0\langle C\rangle \left(\frac{\pi}{2}\right)^{3/2} \frac{\tilde{\omega}_X^2\tilde{z}_X\,\tilde{\omega}_Y^2\tilde{z}_Y}{(\tilde{\omega}_X^2+\tilde{\omega}_Y^2)\sqrt{\tilde{z}_X^2+\tilde{z}_Y^2}} \left(1+\frac{8D\tau}{\tilde{\omega}_X^2+\tilde{\omega}_Y^2}\right)^{-1} \left(1+\frac{8D\tau}{\tilde{z}_X^2+\tilde{z}_Y^2}\right)^{-1/2}. \tag{S39}$$

Eq. S39 now allows us to compute the intensity correlation functions for all photon pairs, which define the FRET correlation function (eq. 2 main text). Eq. 2 (main text) also requires the measured mean FRET value, which will differ from $\varepsilon(t) = 0.5$ due to the different detection volumes for donor and acceptor. It can be computed from the mean photon rates of the two dyes and is given by

$$\langle E\rangle = \frac{\langle n_A(\mathbf{r},t)\rangle}{\langle n_A(\mathbf{r},t)\rangle + \langle n_D(\mathbf{r},t)\rangle} = \frac{\tilde{\omega}_A^2\tilde{z}_A}{\tilde{\omega}_A^2\tilde{z}_A+\tilde{\omega}_D^2\tilde{z}_D}. \tag{S40}$$

With eq. S40, the FRET correlation function that only describes diffusion artifacts is finally given by

$$g_E(\tau) = \tilde{\omega}_A^2 \tilde{z}_A \, \tilde{\omega}_D^2 \tilde{z}_D \langle E \rangle (1 - \langle E \rangle) \frac{\sqrt{2}[G_A(\tau) + G_D(\tau)] - 8G_{AD/DA}(\tau)}{\sqrt{2}[\tilde{\omega}_A^4 \tilde{z}_A^2 G_A(\tau) + \tilde{\omega}_D^4 \tilde{z}_D^2 G_D(\tau)] + 8\tilde{\omega}_A^2 \tilde{z}_A \tilde{\omega}_D^2 \tilde{z}_D G_{AD/DA}(\tau)}$$

with

$$G_A(\tau) = (4D\tau + \tilde{\omega}_A^2)^{-1}(4D\tau + \tilde{z}_A^2)^{-1/2}$$

$$G_D(\tau) = (4D\tau + \tilde{\omega}_D^2)^{-1}(4D\tau + \tilde{z}_D^2)^{-1/2}$$

$$G_{AD/DA}(\tau) = (8D\tau + \tilde{\omega}_A^2 + \tilde{\omega}_D^2)^{-1}(8D\tau + \tilde{z}_A^2 + \tilde{z}_D^2)^{-1/2} \, . \qquad \text{(S41)}$$

Using eq. S39 – 41, we computed the intensity and FRET correlation functions (Supplementary Fig. 2 *middle* and *right*). As a reasonable choice for the size of the excitation volume, we chose $\omega_{ex} = 400\,nm$ and $z_{ex} = 1000\,nm$.[2] The dimensions of the detection volumes for the dye $X$ are then given by $\omega_X = \omega_{ex}\,\lambda_X/\lambda_{ex}$ and $z_X = z_{ex}\,\lambda_X/\lambda_{ex}$. Using eq. S34 we compute the dimensions of the effective detection volumes, which with $\lambda_{ex} = 485\,nm$, $\lambda_D = 520\,nm$, and $\lambda_A = 620\,nm$ result in $\tilde{\omega}_A = 315\,nm$ and $\tilde{z}_A = 788\,nm$ for the acceptor, and $\tilde{\omega}_D = 293\,nm$ and $\tilde{z}_D = 731\,nm$ for the donor. The measured mean FRET efficiency in this case is $\langle E \rangle = 0.555$ instead of the true value 0.5. When we compute the FRET correlation function using eq. S41, we indeed find a decay with a characteristic time of ~630 μs, which is a remnant of the diffusion component in the intensity correlation functions (Supplementary Fig. 1 *middle* and eqs. S39). Yet, the amplitude of this decay is extremely small (~ 0.0013) and undershoots our experimental estimate for the lowest amplitude of ~ 0.002 (Fig. 8b main text).

We therefore conclude that even when accounting for the different effective detection volumes of donor and acceptor, the cancelation of the prominent diffusion decay component present in the intensity correlation functions, is very effective in FRET correlation functions. We would like to note that our theoretical estimate does not consider the burst selection procedure, which effectively crops the integration volume in eqs. S38 and likely diminishes the effect of diffusion even further.

**3. The impact of finite length effects.** It is known that correlation function estimates computed from finite-length trajectories do not only suffer from statistical uncertainties, but they can also be biased. Yet, despite the limited burst length of ~1 ms in diffusion-based smFRET data, we do not observe a severe bias at long lag times in the FRET correlation functions computed with eq.2. To explain the lack of bias, we follow closely the notation of a recent study that investigated finite-length bias in FCS[3]. To this end, and without lack of generality, we consider a signal $X$ that is observed at equally spaced time intervals. We have

$M$ trajectories (bursts), each with $N_i$ time points in the $i^{\text{th}}$ trajectory. The signal at time $j$ in the $i^{\text{th}}$ trajectory is then $X_{ij}$ and given by

$$X_{ij} = \langle X \rangle + \delta X_{ij} \tag{S42}$$

where $\langle X \rangle$ is the true mean value of $X$, and $\delta X_{ij}$ is the deviation from that mean. Notably, $\langle X \rangle$ is not accessible in the experiment. Instead, we estimate this number in our FRET correlation functions as

$$\bar{X} = \frac{1}{\sum_{i=1}^{M} N_i} \sum_{i=1}^{M} \sum_{j=1}^{N_i} X_{ij}. \tag{S43}$$

Instead of eq. S43, there is another possibility to estimate $\langle X \rangle$, namely, by first computing the mean for each trajectory

$$\bar{X}_i = \frac{1}{N_i} \sum_{j=1}^{N_i} X_{ij} \tag{S44}$$

and then to average over all trajectories. Importantly, due to the difference in length of all trajectories, it is clear that this pocedure does not recover eq. S43, i.e.

$$\bar{X} \neq \frac{1}{M} \sum_{i=1}^{M} \bar{X}_i. \tag{S45}$$

With eq. S43, we define the experimental correlation function as

$$\hat{C}(\tau) = \frac{1}{\sum_{i=1}^{M} (N_i - \tau)} \sum_{i=1}^{M} \sum_{j=1}^{N_i - \tau} \left(X_{ij} - \bar{X}\right)\left(X_{ij+\tau} - \bar{X}\right). \tag{S46}$$

To simplify, we define

$$N(\tau) \equiv \sum_{i=1}^{M} (N_i - \tau)\, H(N_i - \tau) \quad \text{with the step function} \quad H(\zeta) = \begin{cases} 0 & \zeta < 0 \\ 1 & \zeta \geq 0 \end{cases}. \tag{S47}$$

When introducing the true mean via eq. S42 and with some rearrangements, the correlation function can be written as

$$\hat{C}(\tau) = \frac{1}{N(\tau)} \sum_{i=1}^{M} \sum_{j=1}^{N_i - \tau} \left(\delta X_{ij} - \frac{1}{N(0)} \sum_{k=1}^{M} \sum_{l=1}^{N_k} \delta X_{kl}\right) \left(\delta X_{ij+\tau} - \frac{1}{N(0)} \sum_{k=1}^{M} \sum_{l=1}^{N_k} \delta X_{kl}\right). \tag{S48}$$

Multiplying the brackets and rearranging the sums gives

$$\hat{C}(\tau) = \frac{1}{N(\tau)} \sum_{i=1}^{M} \sum_{j=1}^{N_i - \tau} \delta X_{ij} \delta X_{ij+\tau} + \left(\frac{1}{N(0)} \sum_{k=1}^{M} \sum_{l=1}^{N_k} \delta X_{kl}\right)^2 - \frac{1}{N(\tau) N(0)} \sum_{i=1}^{M} \sum_{j=1}^{N_i - \tau} \sum_{k=1}^{M} \sum_{l=1}^{N_k} \left(\delta X_{ij} \delta X_{kl} + \delta X_{ij+\tau} \delta X_{kl}\right). \tag{S49}$$

The first term in eq. S49 is the unbiased estimator of the correlation function $C(\tau)$, which differs from the measured function $\hat{C}(\tau)$. The second and third terms are the sources of bias due to the finite length of the trajectories. To determine the bias-terms in $\hat{C}(\tau)$, we average over all possible ensembles, which allows us to estimate the magnitude of the bias. The ensemble average is

$$\langle \hat{C}(\tau) \rangle = \frac{1}{N(\tau)} \sum_{i=1}^{M} \sum_{j=1}^{N_i-\tau} \langle \delta X_{ij} \delta X_{ij+\tau} \rangle + \langle (\bar{X} - \langle X \rangle)^2 \rangle - \frac{1}{N(\tau)N(0)} \sum_{i=1}^{M} \sum_{j=1}^{N_i-\tau} \sum_{k=1}^{M} \sum_{l=1}^{N_k} \left( \langle \delta X_{ij} \delta X_{kl} \rangle + \langle \delta X_{ij+\tau} \delta X_{kl} \rangle \right). \quad \text{(S50)}$$

It is instructive to check the limit in which all trajectories have an identical finite length, i.e., $N_i = N$. In this case, the second term is an estimator of the variance of the mean for which we use the result obtained by Zwanzig[4] (ref. 36 in the main text), which is given by

$$\langle (\bar{X} - \langle X \rangle)^2 \rangle = \frac{2\tau_D}{MN} C(0) \quad \text{(S51)}$$

where $\tau_D$ is the characteristic decay time of the true correlation function. This term is indepent of the time lag $\tau$ and therefore not problematic as it does not distort the shape of the correlation function. However, the third term changes with $\tau$. We can investigate this term better by realizing that the average over a large ensemble allows us to essentially drop the explicit average over different trajectories as indicated by the indices $i$ and $k$. In fact, since the trajectories $i$ and $k$ are uncorrelated when we ignore the recurrence of molecule to the confocal volume, we can express the averages in the third term of eq. S50 as

$$\langle \delta X_{ij} \delta X_{kl} \rangle = \langle \delta X_0 \delta X_{l-j} \rangle \delta_{ik} \quad \text{(S52)}$$

where $\delta_{ik}$ is the Kronecker delta. Hence, only those terms for which $i = k$ will contribute to the sums in the third term in eq. S50, which, after summation over $k$, leads to

$$\langle \hat{C}(\tau) \rangle = C(\tau) + \frac{2\tau_D}{MN} C(0) - \frac{1}{M(N-\tau)MN} \sum_{i=1}^{M} \sum_{j=1}^{N-\tau} \sum_{l=1}^{N} (\langle \delta X_0 \delta X_{l-j} \rangle + \langle \delta X_0 \delta X_{l+\tau-j} \rangle) \quad \text{(S53)}$$

where we used the fact that $N(0) = MN$ and $N(\tau) = M(N - \tau)$ given our assumption of identical length of all trajectories. Eq. S53 does not have a dependence on the index $i$ and hence taking the sum over $i$ finally gives

$$\langle \hat{C}(\tau) \rangle = C(\tau) + \frac{2\tau_D}{MN} C(0) - \frac{1}{MN(N-\tau)} \sum_{j=1}^{N-\tau} \sum_{l=1}^{N} \left( \langle \delta X_0 \delta X_{l-j} \rangle + \langle \delta X_0 \delta X_{l+\tau-j} \rangle \right). \quad \text{(S54)}$$

From eq. S54 it is clear that the lag-time dependent bias term (third term) decays with $1/M$. Hence, for the large number of bursts measured in a diffusion-based smFRET experiment (several thousands), this bias becomes negligible. It is instructive to compare eq. S54 with the result that would be obtained if we first compute the correlation function for each trajectory and then average these correlation functions, which is equivalent to setting $M = 1$ and is therefore given by

$$\langle \hat{C}(\tau) \rangle = C(\tau) + \frac{2\tau_D}{N} C(0) - \frac{1}{N(N-\tau)} \sum_{j=1}^{N-\tau} \sum_{l=1}^{N} \big( \langle \delta X_0 \delta X_{l-j} \rangle + \langle \delta X_0 \delta X_{l+\tau-j} \rangle \big). \qquad \text{(S55)}$$

Clearly, the contribution of the lag-time dependent bias is significantly enhanced in this case.

**4. Coarse-graining of the photophysical model.** To perform simulations of the photophysical model of AlexaFluor488 and AlexFluor594 shown in Fig. 5a in the main text[5], we mapped this scheme with 9 photophysical states onto a coarse-grained (CG) model with 4 CG states. To this end, we combined states with a particular combination of singlet and triplet states. The steady-state populations of the states and kinetic rates in the original model are denoted by $P$ and $k$, respectively, whereas the populations of the CG model are additionally marked by a tilde. The sets of CG states are given by $SS = \{S_0S_0, S_1S_0, S_0S_1, S_1S_1\}$, $TS = \{T_1S_0, T_1S_1\}$, $ST = \{S_1T_0, S_1T_1\}$, and $TT = \{T_1T_1\}$ . The populations of these four states are given by

$$\tilde{P}_{SS} = P_{S_0S_0} + P_{S_1S_0} + P_{S_0S_1} + P_{S_1S_1}$$

$$\tilde{P}_{TS} = P_{T_1S_0} + P_{T_1S_1}$$

$$\tilde{P}_{ST} = P_{S_0T_1} + P_{S_1T_1}$$

$$\tilde{P}_{TT} = P_{T_1T_1} \ . \qquad \text{(S56)}$$

Here, the first subscript indicates the state of the donor and the second subscript indicates the state of the acceptor, $S$ and $T$ indicate singlet and triplet states, respectively, and the numbers indicate ground $(0)$ and first excited state $(1)$. We require that the fluxes between the 9 photophysical states are preserved in the CG model, specifically

$$\tilde{k}_{m\to n}\tilde{P}_m = \sum_{i\in\{m\},j\in\{n\}} k_{i\to j}\, P_i \qquad \text{(S57)}$$

where $\{m\}$ and $\{n\}$ indicate the set of original states that form the coarse-grained states $m$ and $n$ respectively. The CG kinetic rates are given by

$$\tilde{k}_{m\to n} = \frac{\sum_{i\in\{m\},j\in\{n\}} k_{i\to j} P_i}{\sum_{i\in\{m\},j\in\{n\}} P_i}. \qquad \text{(S58)}$$

Where $P_i$ are the steady-state populations in the original model. To parameterize the CG model, we first calculate the dependence of $\tilde{k}_{m\to n}$ as function of the excitation rate $k_{ex} = k_{S_0S_0\to S_1S_0}$. We noted that the rates $\tilde{k}_{m\to n}$ for the transitions $SS \to TS$, $SS \to ST$, $ST \to TT$, and $TS \to TT$ depend approximately linearly on $k_{ex}$ (Supplmentary Fig. 2 *top*). To account for this dependence, we assumed a linear dependence of these rates in the parametrization of the CG model

$$\tilde{k}_{m\to n} = \tilde{a}_{m\to n} k_{ex}. \tag{S59}$$

On the other hand, transitions in the opposite direction, i.e., $TS \to SS$, $ST \to SS$, $TT \to ST$, and $TT \to TS$, only show a weak or no dependence on $k_{ex}$ (Supplmentary Fig. 2 *bottom*) and are therefore described by constants

$$\tilde{k}_{n\to m} = \tilde{b}_{n\to m}. \tag{S60}$$

The set of $\tilde{a}_{m\to n}$ and $\tilde{b}_{n\to m}$ constitute 8 fitting parameters that were determined by fitting eq. S59-60 to the $k_{ex}$-dependence of $\tilde{k}_{m\to n}$ and $\tilde{k}_{n\to m}$ that were computed from the full 9-state model via eq. S58. Each of the CG states is also associated with 2 emission rates, one for the emission of donor photons and the other for the emission of acceptor photons. The emission rates are given by

$$\tilde{n}_{SS}^{A}(k_{ex}) = \frac{k_{S_0S_1\to S_0S_0} P_{S_0S_1}(k_{ex}) + k_{S_1S_1\to S_1S_0} P_{S_1S_1}(k_{ex})}{\sum_{i\in SS} P_i(k_{ex})}$$

$$\tilde{n}_{SS}^{D}(k_{ex}) = \frac{k_{S_1S_0\to S_0S_0} P_{S_1S_0}(k_{ex}) + k_{S_1S_1\to S_0S_1} P_{S_1S_1}(k_{ex})}{\sum_{i\in SS} P_i(k_{ex})}$$

$$\tilde{n}_{TS}^{A}(k_{ex}) = \frac{k_{T_1S_1\to T_1S_0} P_{T_1S_1}(k_{ex})}{\sum_{i\in TS} P_i(k_{ex})}$$

$$\tilde{n}_{TS}^{D}(k_{ex}) = 0$$

$$\tilde{n}_{ST}^{A}(k_{ex}) = 0$$

$$\tilde{n}_{ST}^{D}(k_{ex}) = \frac{k_{S_1T_1\to S_0T_1} P_{S_1T_1}(k_{ex})}{\sum_{i\in ST} P_i(k_{ex})}$$

$$\tilde{n}_{TT}^{A}(k_{ex}) = \tilde{n}_{TT}^{D}(k_{ex}) = 0\ . \tag{S61}$$

We again computed these emission rates as function of $k_{ex}$ and used a linear relationship of the form $\tilde{n}_l^X = \tilde{m}_l^X\, k_{ex}$ where $X$ and $l$ indicate the photon color and CG state, respectively. With these definitions, the dynamics of the CG model can be written in matrix notation as

$$\frac{d\tilde{\mathbf{p}}}{dt} = \tilde{\mathbf{K}}\,\tilde{\mathbf{p}} \tag{S62}$$

with the state vector

$$\tilde{\mathbf{p}} = (\tilde{P}_{SS} \quad \tilde{P}_{TS} \quad \tilde{P}_{ST} \quad \tilde{P}_{TT})^{\mathrm{T}} \tag{S63}$$

and the rate matrix $\tilde{\mathbf{K}} = \tilde{\mathbf{K}}_0 + \tilde{\mathbf{K}}_1$ with

$$\tilde{\mathbf{K}}_0 = \begin{pmatrix} 0 & \tilde{b}_{TS\to SS} & \tilde{b}_{ST\to SS} & 0 \\ 0 & -\tilde{b}_{TS\to SS} & 0 & \tilde{b}_{TT\to TS} \\ 0 & 0 & -\tilde{b}_{ST\to SS} & \tilde{b}_{TT\to SS} \\ 0 & 0 & 0 & -(\tilde{b}_{TT\to TS} + \tilde{b}_{TT\to SS}) \end{pmatrix}$$

$$\tilde{\mathbf{K}}_1 = k_{ex} \begin{pmatrix} -(\tilde{a}_{SS\to TS} + \tilde{a}_{SS\to ST}) & 0 & 0 & 0 \\ \tilde{a}_{SS\to TS} & -\tilde{a}_{TS\to TT} & 0 & 0 \\ \tilde{a}_{SS\to ST} & 0 & -\tilde{a}_{ST\to TT} & 0 \\ 0 & \tilde{a}_{TS\to TT} & \tilde{a}_{ST\to TT} & 0 \end{pmatrix}. \tag{S64}$$

Here, $\tilde{\mathbf{K}}_0$ contains all $T \to S$ transitions that are independent of the excitation rate and $\tilde{\mathbf{K}}_1$ contains all excitation-dependent $S \to T$ transitions. The emission rate matrices for acceptor ($\tilde{\mathbf{V}}_A$) and donor ($\tilde{\mathbf{V}}_D$) are given by

$$\tilde{\mathbf{V}}_A = k_{ex} \begin{pmatrix} \tilde{m}_{SS}^A & 0 & 0 & 0 \\ 0 & \tilde{m}_{TS}^A & 0 & 0 \\ 0 & 0 & 0 & 0 \\ 0 & 0 & 0 & 0 \end{pmatrix} \text{ and } \tilde{\mathbf{V}}_D = k_{ex} \begin{pmatrix} \tilde{m}_{SS}^D & 0 & 0 & 0 \\ 0 & 0 & 0 & 0 \\ 0 & 0 & \tilde{m}_{ST}^D & 0 \\ 0 & 0 & 0 & 0 \end{pmatrix}. \tag{S65}$$

The rates for the full photophysical model and the value of the CG version can be found in Supplementary Table 1 and 2.

**Supplementary Table 1.** Parameters of the 9-state photophysical model of the dye pair AlexaFluor488 (A) and AlexaFluor594 (D) determined by Nettels et al. (ref.[5]).

| Parameter (Fig. 5a main text) | Parameter (Nettels *et al.*) | Value |
|---|---|---|
| $\boldsymbol{k_{S_0S_0\to S_1S_0} = k_{S_0S_2\to S_1S_1} = k_{S_0T_1\to S_1T_1}}$ | $k_{ex}$ | $0-0.08\,ns^{-1}$ |
| $\boldsymbol{k_{S_1S_0\to S_0S_1}}$ | $k_T$ | $0.25\,(R_0/r)^6\,ns^{-1}$ |
| $\boldsymbol{k_{S_1S_0\to S_0S_0} = k_{S_1S_1\to S_0S_1} = k_{S_1T_1\to S_0T_1}}$ | $k_D$ | $0.25\,ns^{-1}$ |
| $\boldsymbol{k_{S_0S_1\to S_0S_0} = k_{S_1S_1\to S_1S_0} = k_{T_1S_1\to T_1S_0}}$ | $k_A$ | $0.25\,ns^{-1}$ |
| $\boldsymbol{k_{S_1S_1\to S_1T_1} = k_{S_0S_1\to S_0T_1} = k_{T_1S_1\to T_1T_1}}$ | $k_{S_1\to T_1}$ (A) | $10^{-3}\,ns^{-1}$ |
| $\boldsymbol{k_{S_1T_1\to S_1S_0} = k_{T_1T_1\to T_1S_0} = k_{S_0T_1\to S_0S_0}}$ | $k_{T_1\to S_0}$ (A) | $3\,10^{-4}\,ns^{-1}$ |
| $\boldsymbol{k_{S_1S_0\to T_1S_0} = k_{S_1S_1\to T_1S_1} = k_{S_1T_1\to T_1T_1}}$ | $k_{S_1\to T_1}$ (D) | $10^{-3}\,ns^{-1}$ |
| $\boldsymbol{k_{T_1S_1\to S_0S_1} = k_{T_1T_1\to S_0T_1} = k_{T_1S_0\to S_0S_0}}$ | $k_{T_1\to S_0}$ (D) | $3\,10^{-4}\,ns^{-1}$ |
| $\boldsymbol{k_{S_1S_1\to S_0S_1}}$ | $k_{SSA}$ | $k_T$ |
| $\boldsymbol{k_{S_1T_1\to S_0T_1}}$ | $k_{STA}$ | $3\,k_T$ |
| $\boldsymbol{k_{S_0S_0\to S_0S_1} = k_{S_1S_0\to S_1S_1} = k_{T_1S_0\to T_1S_1}}$ | $\alpha k_{ex}$ | $0.05\,k_{ex}$ |
| $\boldsymbol{R_0}$ | $R_0$ | $5.4\,nm$ |

**Supplementary Table 2.** Parameters of the CG model of the dye pair AlexaFluor488 (A) and AlexaFluor594 (D).

| Parameter | Value |
|---|---|
| $\boldsymbol{\tilde{a}_{SS\to TS}}$ | $1.8\,10^{-3}ns^{-1}$ |
| $\boldsymbol{\tilde{a}_{SS\to ST}}$ | $1.84\,10^{-3}ns^{-1}$ |
| $\boldsymbol{\tilde{a}_{TS\to TT}}$ | $2.0\,10^{-4}ns^{-1}$ |
| $\boldsymbol{\tilde{a}_{ST\to TT}}$ | $9.5\,10^{-4}ns^{-1}$ |
| $\boldsymbol{\tilde{b}_{TS\to SS}}$ | $3.0\,10^{-4}ns^{-1}$ |
| $\boldsymbol{\tilde{b}_{ST\to SS}}$ | $3.0\,10^{-4}ns^{-1}$ |
| $\boldsymbol{\tilde{b}_{TT\to TS}}$ | $3.0\,10^{-4}ns^{-1}$ |
| $\boldsymbol{\tilde{b}_{TT\to ST}}$ | $3.0\,10^{-4}ns^{-1}$ |
| $\boldsymbol{\tilde{m}^A_{SS}}$ | 0.487 |
| $\boldsymbol{\tilde{m}^D_{SS}}$ | 0.465 |
| $\boldsymbol{\tilde{m}^A_{TS}}$ | 0.05 |
| $\boldsymbol{\tilde{m}^D_{TS}}$ | 0.0 |
| $\boldsymbol{\tilde{m}^A_{ST}}$ | 0.0 |
| $\boldsymbol{\tilde{m}^D_{ST}}$ | 0.24 |
| $\boldsymbol{\tilde{m}^A_{TT}}$ | 0.0 |
| $\boldsymbol{\tilde{m}^D_{TT}}$ | 0.0 |

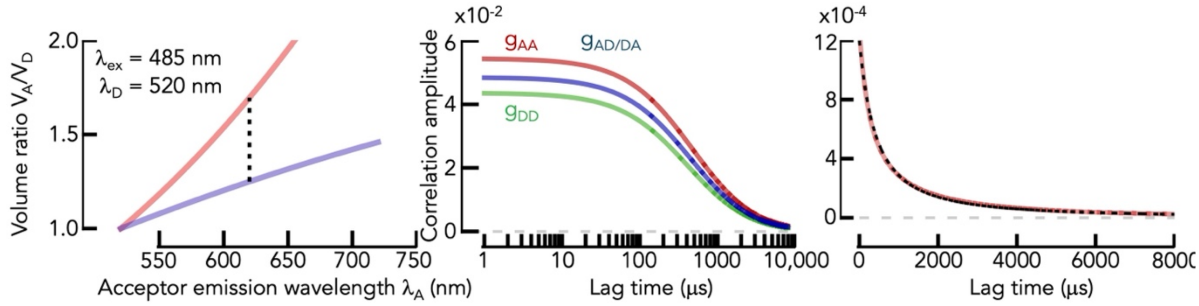

**Supplementary Fig. 1. Impact of different apparent detection profiles on the suppression of the diffusion component.** (*Left*) Ratio of the emission profiles (red) and apparent detection profiles (blue) of acceptor and donor as function of the emission wavelength of the acceptor as computed with eq. S35. (*Middle*) Diffusion component of the acceptor and donor autocorrelation and cross-correlation functions of a particle with $\varepsilon(t) = 0.5$ calculated with eq. S39. (*Right*) FRET correlation function computed for the process shown in Middle.

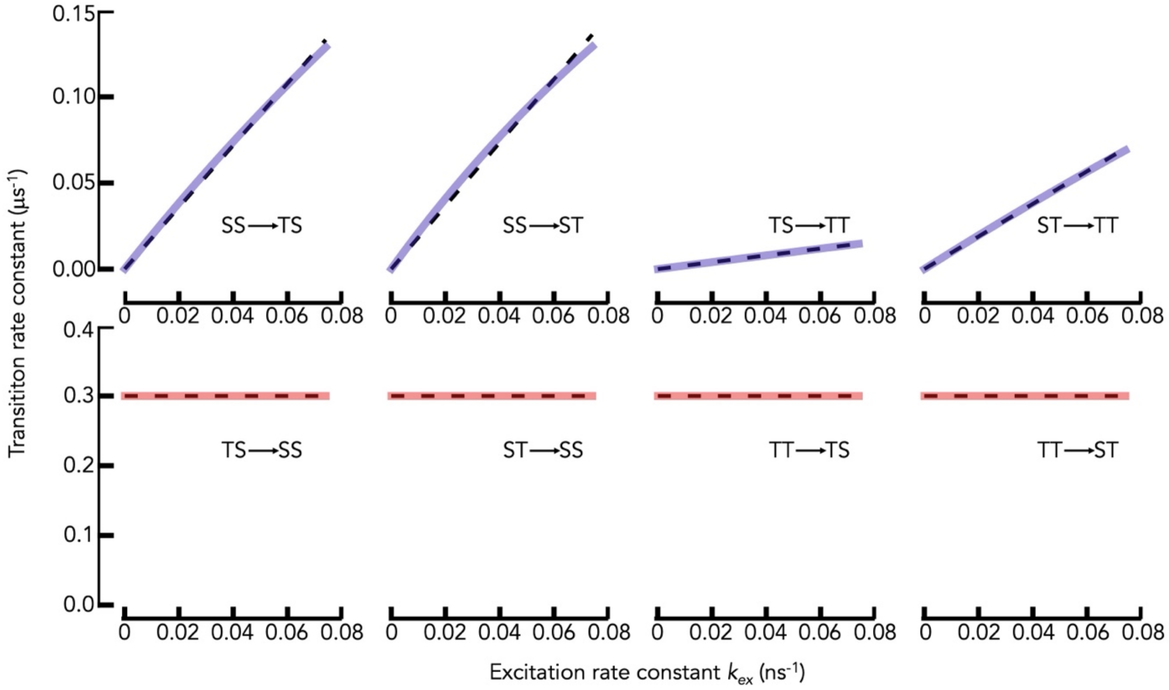

**Supplementary Fig. 2. Transition rates of the CG model.** Solid lines indicate the CG rates calculated from the full 9-state model using eq. S58. The transition type is indicated. Black dashed lines are fits with eq. S59-60.

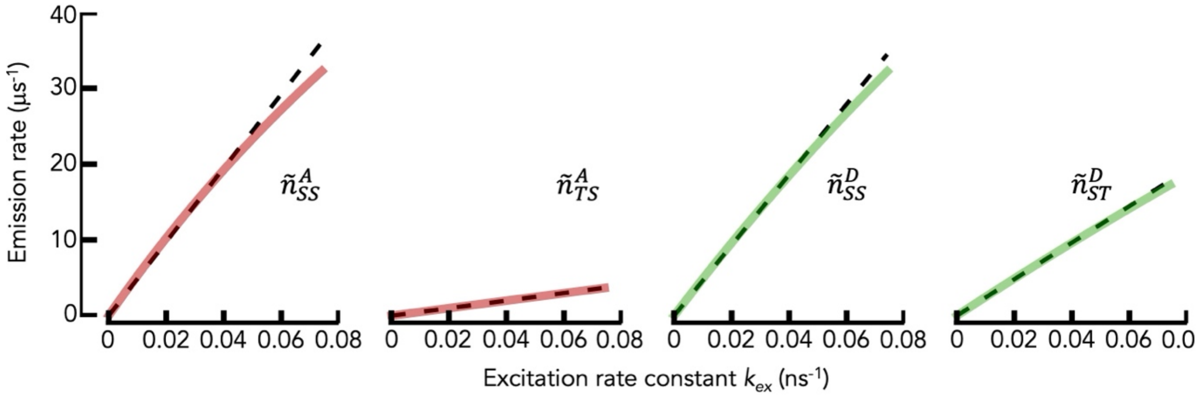

**Supplementary Fig. 3. Emission rates of the CG model.** Solid lines indicate the CG rates calculated from the full 9-state model using eq. S61. The emission type is indicated. Black dashed lines are linear fits.

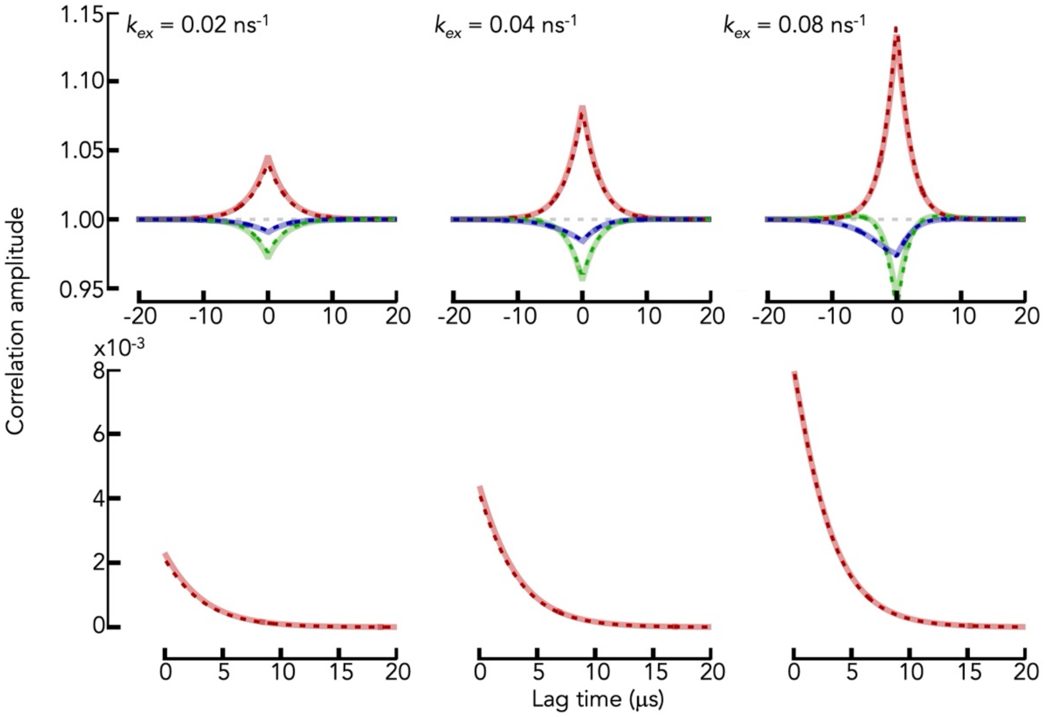

**Supplementary Fig. 4. Benchmarking the CG model against the 9-state model.** Correlation ratios (top) and FRET correlation functions (bottom) were computed from the 9-state model (solid lines) and from the parametrized CG model (dashed lines). The color code in the top panel indicates the normalized correlation ratios $f_{AA}$ (red), $f_{DD}$ (green), and $f_{AD/DA}$ (blue). From left to right, the excitation rate increases (indicated).
